# Independent assessment and improvement of wheat genome assemblies using Fosill jumping libraries

**DOI:** 10.1101/219352

**Authors:** Fu-Hao Lu, Neil McKenzie, George Kettleborough, Darren Heavens, Matthew D. Clark, Michael W. Bevan

**Author notes:** Joint first Authors (Fu-Hao Lu, Neil McKenzie).

## Abstract

**Background:** The accurate sequencing and assembly of very large, often polyploid, genomes remain a challenging task, limiting long range sequence information and phased sequence variation for applications such as plant breeding. The 15 Gb hexaploid bread wheat genome has been particularly challenging to sequence, and several contending approaches recently generated accurate long range assemblies. Understanding errors in these assemblies is important for optimising future sequencing and assembly approaches and for comparative genomics.

**Results:** Here we use a Fosill 38 Kb jumping library to assess medium and longer range order of different publicly available wheat genome assemblies. Modifications to the Fosill protocol generated longer Illumina sequences and enabled comprehensive genome coverage. Analyses of two independent BAC based chromosome-scale assemblies, two independent Illumina whole genome shotgun assemblies, and a hybrid long read (PacBio) and short read (Illumina) assembly were carried out. We revealed a variety of discrepancies using Fosill mate-pair mapping and validated several of each class. In addition, Fosill mate-pairs were used to scaffold a whole genome Illumina assembly, leading to a three-fold increase in N50 values.

**Conclusions:** Our analyses, using an independent means to validate different wheat genome assemblies, show that whole genome shotgun assemblies are significantly more accurate by all measures compared to BAC-based chromosome scale assemblies. Although current whole genome assemblies are reasonably accurate and useful, additional steps will be needed for the rapid, cost effective and complete sequencing and assembly of wheat genomes.

## Background

Genome sequence assemblies are key foundations for many biological studies, therefore the accuracy of sequence assemblies and their long-range order is a fundamental prerequisite for their use. Multiple types of differences in the information content of DNA molecules, from single nucleotide polymorphisms (SNPs) to large-scale structural variation (SV), form part of natural genetic variation that can cause phenotypic variation [1,2]. Distinguishing such *bona fide* variation from apparent variation generated by sequence and assembly methods is therefore a critically important activity in genomics.

Sequence assemblies are generally incomplete and contain multiple types of errors, reducing their information content. Gaps in assemblies can occur where no sequence reads were generated for that region, but this is now increasingly unlikely given the very deep coverage achievable by short read sequencing, improved sequence chemistry, and template preparation methods that avoids bias, such as that introduced by PCR [3]. Closely related repetitive DNA sequences can lead to incorrect joins in assemblies, or to an unresolvable assembly graph that breaks an assembly. Assemblies can be either joined or broken inadvertently by closely related or polymorphic sequences that cause alternate, multiple, or collapsed assemblies, for example in assemblies of polyploid organisms [4]. Errors and incompleteness can obscure important genomic information such as the correct order (phasing) of sequence variants.

A broad spectrum of sequence and assembly artefacts can be distinguished from natural sequence variation, structural variants identified, and sequence variation phased, using long-range sequence information. Fosmid clones, which have large precisely-sized inserts due to lambda phage packaging, have been sequenced to close gaps in human genome assemblies [5] and to establish longer-range sequence haplotypes [6,7]. Earlier uses of fosmid clones for bulk sequencing [8,9] were supplanted by sequencing libraries of larger insert Bacterial Artificial Chromosome (BAC) clones [10]. Sequences of long single molecules generated by PacBio Single Molecule Real Time (SMRT) and Nanopore technologies are increasingly used for defining long-range gene order and for *de novo* genome assembly [11,12]. Linked read technologies such as 10X Genomics reads are also beginning to be widely used for long-range ordering of scaffolds assembled from short reads, and for identifying structural variation [13]. SMRT is also often used in hybrid approaches that utilise Illumina assemblies to improve the accuracy of single molecule reads [14]. Thus, there are several approaches available for generating and assessing the long-range integrity of genome assemblies.

These improvements in sequence read length and assembly procedures are enabling the creation of genomic resources for even the largest genomes. These include the genomes of grasses and gymnosperm trees, which have massive repetitive DNA tracts comprising about 80% of their genomes. The 22 Gb genome of loblolly pine (*Pinus taeda*), initially assembled from Illumina paired end sequence reads [15], has been significantly improved using SMRT sequencing [16]. 10X Genomics linked reads were used to generate an eight-fold increase in scaffold NG50 sizes of sugar pine (*P. lambertiana*) genome assemblies to nearly 2 Mb [17]. Bread wheat (*Triticum aestivum*) has a large 15 Gb allohexaploid genome consisting of three closely-related and separately maintained A, B and D genomes [18]. Assemblies of Illumina whole genome shotgun sequences were assigned to their correct genome [19], but the assembly was still highly fragmented. A near-complete and highly contiguous assembly of Illumina paired-end and mate-pair reads from wild emmer wheat (*Triticum turgidum*), a tetraploid progenitor of bread wheat has also recently been published [20]. Finally, long SMRT sequence reads integrated with Illumina sequence coverage increased the size and contiguity of maize [21], a diploid wheat progenitor [14] and hexaploid wheat genome assemblies [22]. It is not known how these assemblies differ in error types which may obscure true genetic variation.

Generating and assessing accurate long-range genome information from the large genomes of crop plants is necessary for identifying haplotypes used by breeders and for mapping large-scale structural variation contributing to agronomic performance [23]. Therefore, assessing the fidelity of longer-range genome assemblies is important for their applications to crop improvement. Here we use mate-paired sequences of wheat fosmid clones to assess three different wheat whole genome assemblies and two BAC-based wheat chromosome assemblies. Our analyses have identified a range of assembly issues and may help to identify optimal approaches to wheat genome assembly. Integrating fosmid end-sequences into scaffolds also increased scaffold sizes of both fragmentary and more contiguous assemblies.

## Results

### Creating and assessing a wheat fosmid clone library

Fosmid clone libraries have been used to assess genome assemblies and identify structural variation in human [24,25] and pine genomes [16]. Fosmids are used because DNA is cloned in a precise range of 38±3 Kb by efficient packaging of phage lambda and cohesive end circularisation. Fosmid clone inserts have been converted to Illumina sequencing templates to generate 38 Kb mate-pair “jumping libraries” for improving the assemblies of the mouse genome [26]. In genomes with extensive tracts of closely-related repeats that have been challenging to assemble, fosmid jumping libraries could provide an independent means to both assess the fidelity of assemblies and to improve them. Several different hexaploid and tetraploid wheat chromosome and whole genome assemblies have been generated using different approaches [19,20,22,27], and assessing these could provide information needed to identify optimal approaches to wheat genome assembly.

To explore the potential of fosmid jumping libraries for assessing and improving wheat genome assemblies, we first carried out a simulation, using 38 Kb mate-pairs, of whole genome shotgun assemblies of three long 3.5 - 4.1 Mb scaffolds of wheat chromosome 3B generated by sequencing a manually curated physical map of BACs [27]. Simulation settings used different paired-end distances, read lengths and sequence coverage on the chromosome 3B scaffolds to assess how 38 Kb mate-paired reads, read-depth and read-length contributed to re-assembly (Additional File 1). Addition of 38 Kb mate-pair reads was required for accurate and complete reconstruction of all three scaffolds under these conditions. Paired-end read lengths between 100 - 250 bp were then assessed using a common combination of mate pair distances and sequence coverage. Reads of over 200 bp were required for consistent reassembly of all three scaffolds. Finally, simulation of sequence coverage of 38 kb mate pair reads of length 250 bp showed that consistent re-assembly of all three scaffolds required sequence coverage of approximately 0.75× (Additional File 1). Taken together, these simulations showed that 38 Kb paired-end 250 bp reads with a sequence coverage of approximately 0.75× (>50x physical coverage) could be used to guide and assess assemblies of the wheat genome.

The Fosill vector system was developed for converting fosmid clones to Illumina paired-end read templates [28]. We modified this Fosill conversion protocol to generate long paired-end 250 bp Illumina reads, to maximise library complexity, and to minimise clonal- and PCR-based amplification bias. Both of these modifications were required to maximise unique matches of paired-end reads to the highly repetitive polyploid wheat genome, and to maximise sequence coverage of the large genome. Additional File 2 describes the modified protocols for library preparation and paired-end read analyses. These involved increasing the time of nick-translation to between 50-60 minutes on ice to generate inverse PCR products with a peak size distribution of 785-860 bp (Additional File 2). This minimised overlap of 250bp reads from either end of the PCR product. For each pool of 5-10M Fosill clones, a small sample of the circularised template was amplified for up to 16 cycles, and the minimum number of cycles required (generally 12-13) to generate sufficient template for sequencing was estimated.

Table 1 in Additional File 2 summarises the Fosill libraries produced and the paired-end sequences generated from them. Paired-end reads that overlapped each other on the template were discarded (2.61%), while 11.91% of the raw reads were excluded after vector/adapter sequence and quality trimming. The final number of 576 M paired-end sequences (85.5% of the total reads) were generated from 54.61 M Fosill clones (124× physical coverage). These were then mapped to the chromosome 3B pseudomolecule to measure the insert size distribution of the libraries (Additional File 3). Figure 1A shows the size distribution of 588,268 mapped read pairs, which had a mean estimated insert size of approximately 37,725 bp. This is the expected insert size range in the Fosill4 vector [28], and demonstrated successful size selection during packaging. Figure 1B shows the distribution of mate-pairs mapped in 100 kb windows across chromosome 3B BAC pseudomolecule. Reads with a depth of ≤5 covered 494 Mb of the total 833 Mb chromosome, accounting for 59% of the chromosome sequence. Their even distribution across the pseudomolecule indicated that the libraries were representative of the entire chromosome. There were approximately 30 distinct peaks of greatly increased read-depth (Figure 1B) in the 100 kb windows across chromosome 3B. These probably correspond to mate-pairs spanning approximately 40 Kb repeated regions common to multiple genomic loci. These reads accounts for 80% of the alignments but covered only 4.3% of chromosome 3B. For all subsequent analyses only Fosill mate-pairs of sequence depth ≤5 were used. Finally, reads that mapped to multiple locations, which lacked a paired read in the expected genomic location, or which had a paired read in the incorrect orientation, were removed.

**Figure 1.**
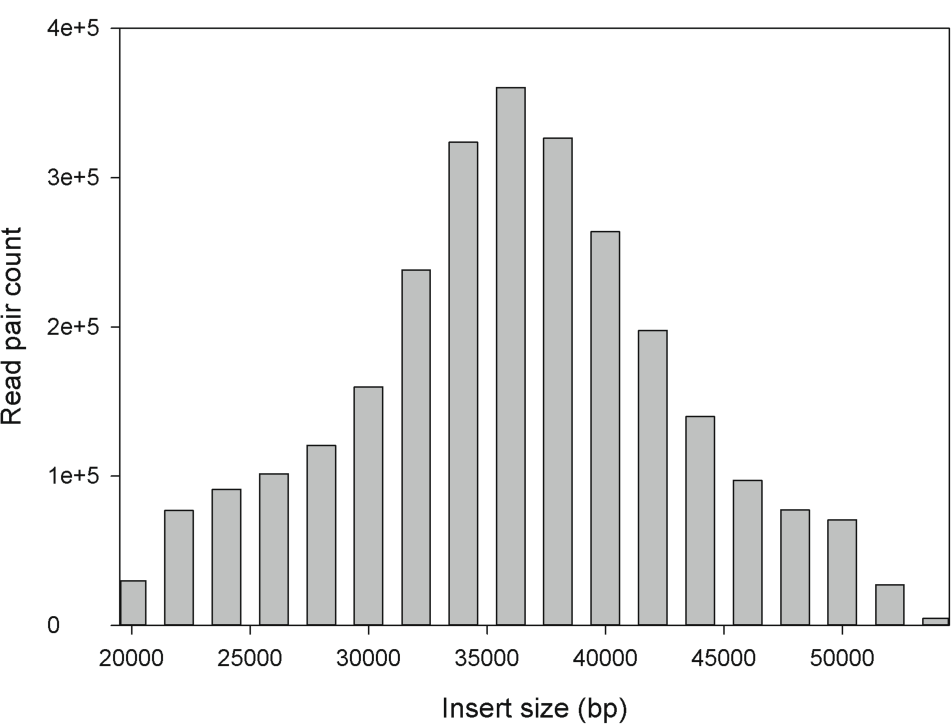
Determination of Fosill mate-pair distance distributions on chromosome 3B. **A**. 576 M quality controlled paired-end sequences were mapped to the chromosome 3B pseudomolecule. 588,268 read pairs were mapped and the insert sizes calculated. The mean insert size was 37,725 Kb.

**Figure B.**
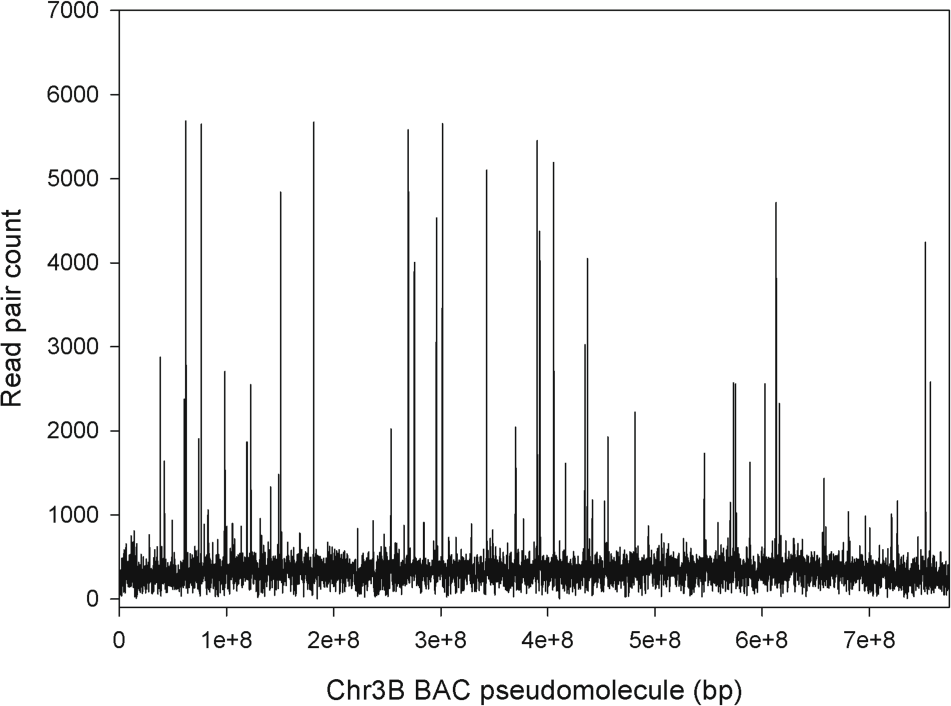
Fosill mate-pairs were mapped in 100 Kb bins along chromosome 3B to assess the depth and evenness of coverage. Coverage was generally even across the entire chromosome, with approximately 30 very high copy peaks that are probably due to Fosill mate-pairs from highly related 40 Kb+ regions from across the genome. Most mate-pairs mapped to a depth of ≤5 and were used for subsequent analyses.

### Using Fosill mate pairs to assess wheat chromosome sequence assemblies

The even representation of long mate-paired reads across the chromosome 3B pseudomolecule indicated their suitability for assessing wheat sequence assemblies and for making new joins in wheat sequence scaffolds. For assessing assemblies, a windows-based filter was developed to identify sets of ≥5 unique neighbouring Fosill sequence reads in a “driver” window of <10 kb and their ≥5 mate-pair reads in a “follower” window of <20 kb on chromosome and genome assemblies. The vast proportion of mate-paired reads fell within this distance distribution (Additional File 3, Figure 1). Using this approach to map Fosill reads, we aimed to identify different types of paired-end matches to genome sequence assemblies. These can be used to identify genome assemblies consistent with the 37.7 kb mate-pair distances +/− sd, to identify possible new joins between assemblies, and to identify different types of inconsistencies in the range of current publicly available wheat genome assemblies. Figure 2A illustrates the possible types of Fosill paired-end matches to assemblies.

**Figure 2.**
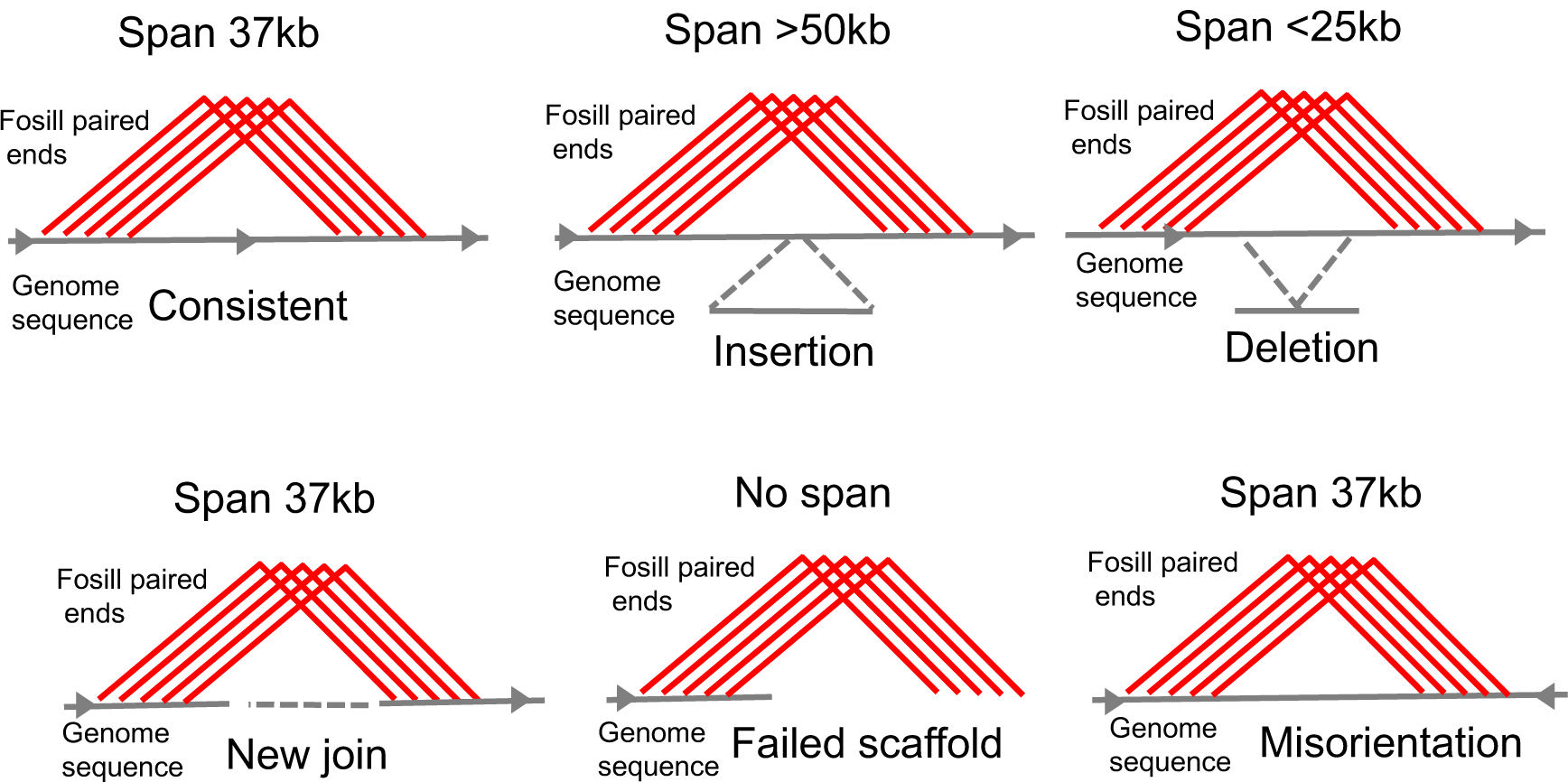
Using Fosill mate-pair matches to identify discrepancies in wheat chromosome and genome assemblies. **A**. The schematic describes different classes of matches of Fosill mate-pair sequences to wheat chromosome and genome assemblies. Consistent assemblies matched a span of ≥5 mate-pairs in a sliding 10 Kb “driver” window that matched their mate in a 20 Kb “follower” window at a distance of 37 Kb +/− sd in the correct orientation. Where mate-pairs spanned more than 50 Kb (approximately 3 sd) this was construed to be due to an aberrant insertion in the underlying assembly. Spans <25 Kb (approximately 3 sd) were construed to be due to an aberrant deletion in the assembly. Mis-orientations of the mate-pairs indicated a mis-oriented assembly, and no span a mis-join in the assembly. New joins were also identified. Drawing not to scale.

Tables 1A-1E show the outcomes of mapping Fosill paired-end reads to BAC-based bread wheat chromosome assemblies of chromosome 3B [27], TGACv1 Illumina assemblies of 3B [19], the Triticum 3.0 whole genome assembly of Pacbio SMRT and Illumina sequences of chromosomes 3B and 3DL [22], and DeNovoMagic assemblies of Illumina sequences from wild emmer wheat (WEW) chromosome 3B [20]. We also assessed an assembly of hexaploid wheat chromosome 3DL from sequenced BACs in a minimal tiling path using an automated pipeline (Additional File 4). A set of larger whole genome assemblies of the TGACv1 Illumina wheat genome were also assessed. These assemblies represent diverse approaches to sequencing wheat chromosomes and chromosome arms, including manually curated and automated BAC-based assemblies, two different Illumina-based assembly methods, and a combined Illumina and Pacific Biosciences SMRT assembly of wheat chromosomes.

**Table 1A.**
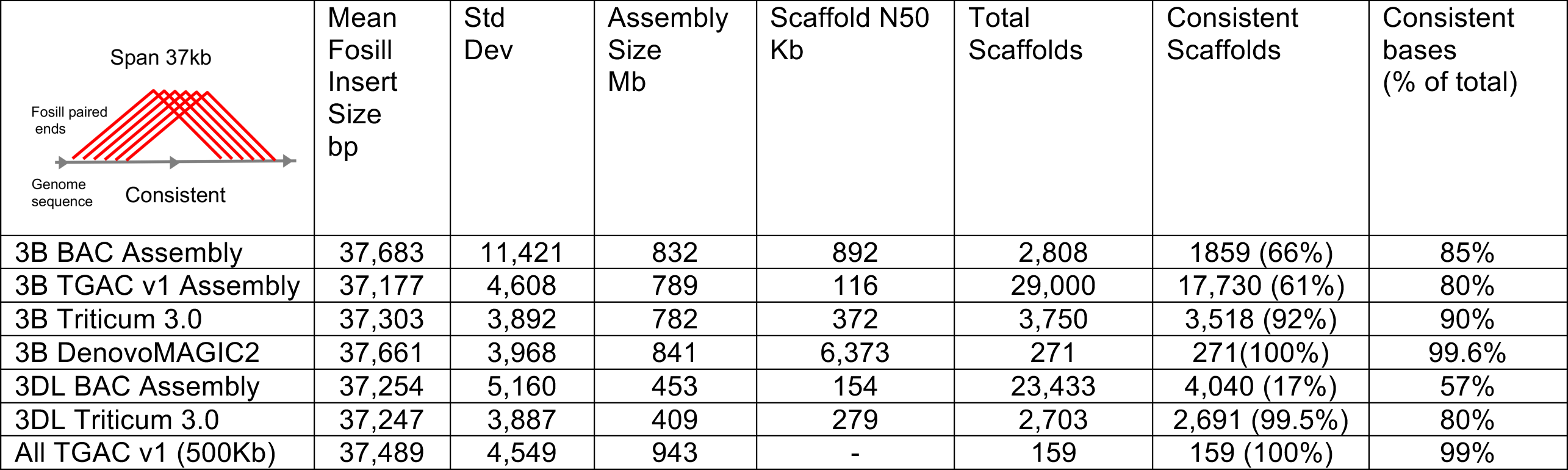
Summary of Fosill mate-pair alignments to different publically-available assemblies of chromosome 3B and 3DL, and the TGACv1 whole genome assembly. The consistency of mapping is shown according to the number of assemblies with consistent matches, and the percentage of bases included in consistent matches to Fosill mate-pairs.

| Span 37kb<br>Fosill paired ends<br>Genome sequence<br>Consistent | Mean Fosill Insert Size bp | Std Dev | Assembly Size Mb | Scaffold N50 Kb | Total Scaffolds | Consistent Scaffolds | Consistent bases (% of total) |
| --- | --- | --- | --- | --- | --- | --- | --- |
| 3B BAC Assembly | 37,683 | 11,421 | 832 | 892 | 2,808 | 1859 (66%) | 85% |
| 3B TGAC v1 Assembly | 37,177 | 4,608 | 789 | 116 | 29,000 | 17,730 (61%) | 80% |
| 3B Triticum 3.0 | 37,303 | 3,892 | 782 | 372 | 3,750 | 3,518 (92%) | 90% |
| 3B DenovoMAGIC2 | 37,661 | 3,968 | 841 | 6,373 | 271 | 271 (100%) | 99.6% |
| 3DL BAC Assembly | 37,254 | 5,160 | 453 | 154 | 23,433 | 4,040 (17%) | 57% |
| 3DL Triticum 3.0 | 37,247 | 3,887 | 409 | 279 | 2,703 | 2,691 (99.5%) | 80% |
| All TGAC v1 (500Kb) | 37,489 | 4,549 | 943 | - | 159 | 159 (100%) | 99% |

**Table 1B.**
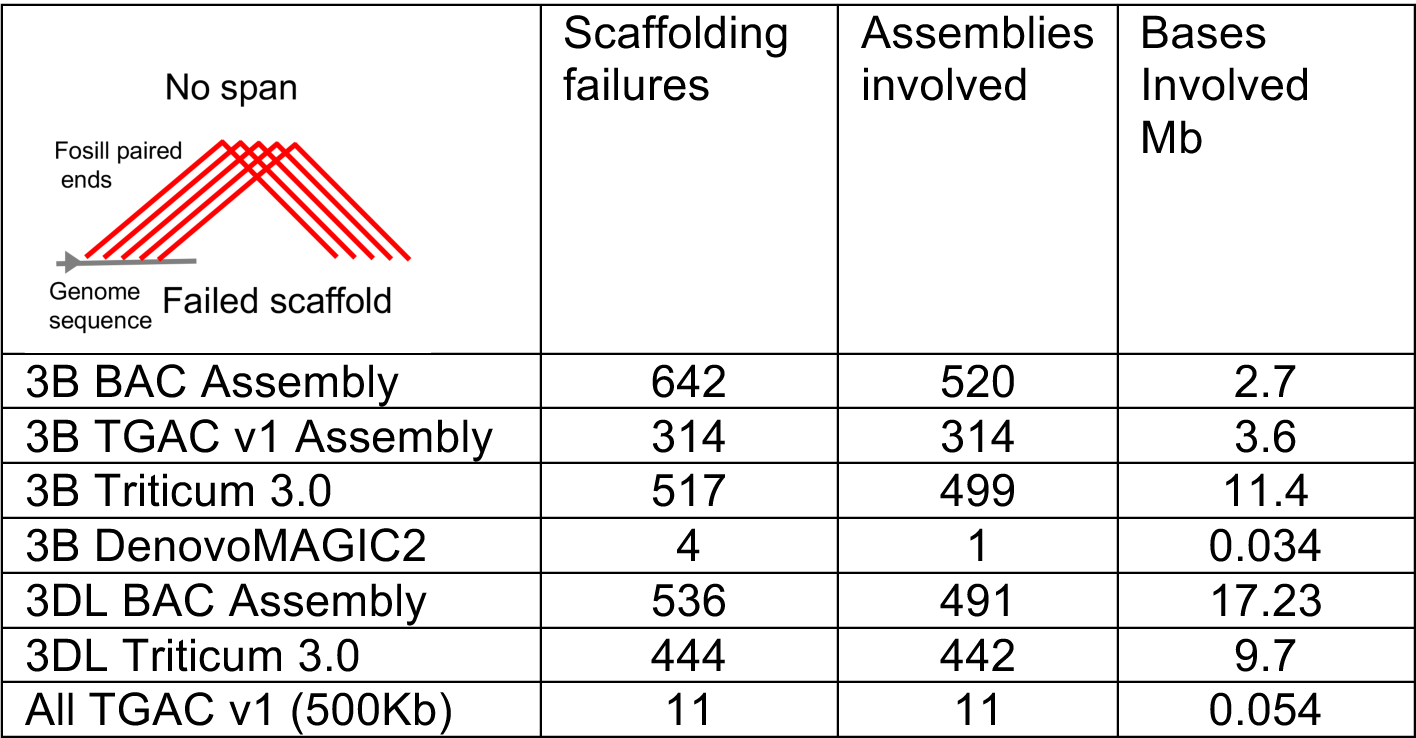
Potential failed scaffolding.

|  |  |  |  |
| --- | --- | --- | --- |
| No span | Scaffolding failures | Assemblies involved | Bases Involved Mb |
| 3B BAC Assembly | 642 | 520 | 2.7 |
| 3B TGAC v1 Assembly | 314 | 314 | 3.6 |
| 3B Triticum 3.0 | 517 | 499 | 11.4 |
| 3B DenovoMAGIC2 | 4 | 1 | 0.034 |
| 3DL BAC Assembly | 536 | 491 | 17.23 |
| 3DL Triticum 3.0 | 444 | 442 | 9.7 |
| All TGAC v1 (500Kb) | 11 | 11 | 0.054 |

**Table 1C.**
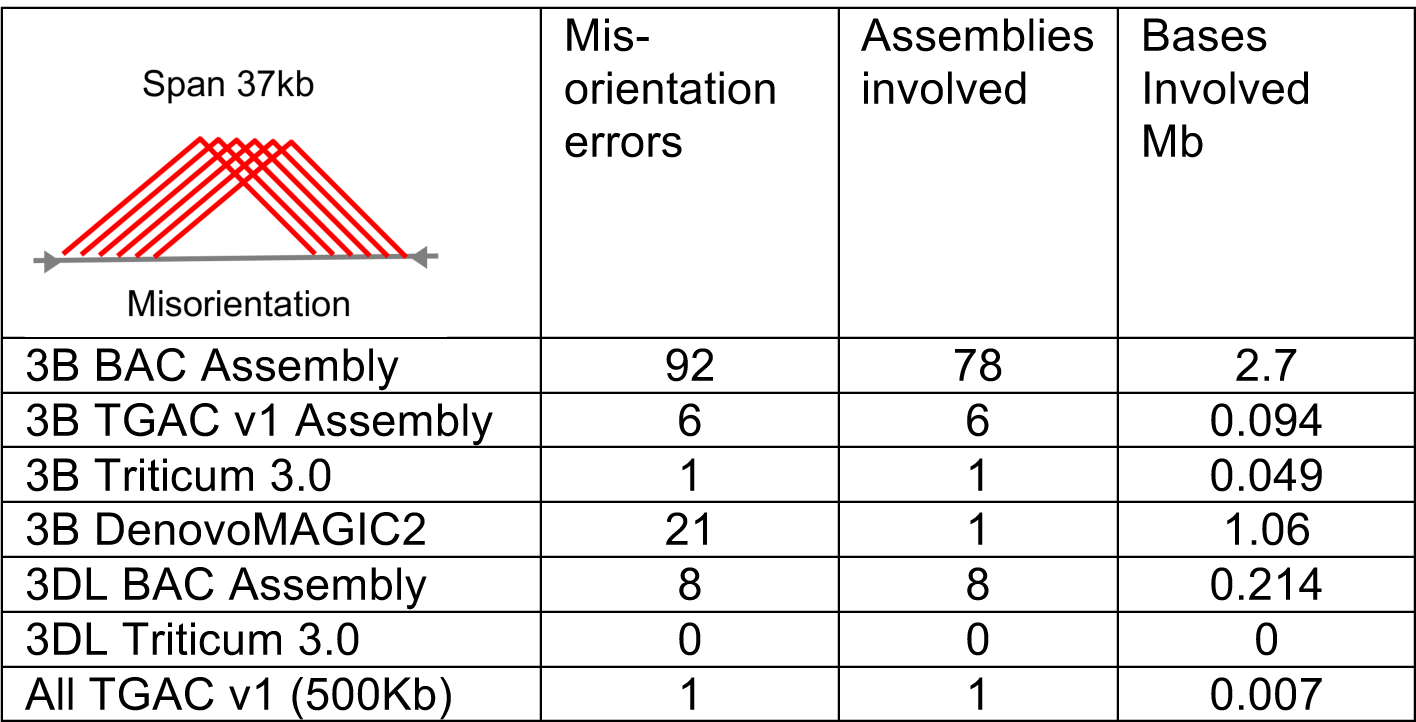
Potential mis-orientations in scaffolds.

|  |  |  |  |
| --- | --- | --- | --- |
| Span 37kb | Mis-orientation errors | Assemblies involved | Bases Involved Mb |
| 3B BAC Assembly | 92 | 78 | 2.7 |
| 3B TGAC v1 Assembly | 6 | 6 | 0.094 |
| 3B Triticum 3.0 | 1 | 1 | 0.049 |
| 3B DenovoMAGIC2 | 21 | 1 | 1.06 |
| 3DL BAC Assembly | 8 | 8 | 0.214 |
| 3DL Triticum 3.0 | 0 | 0 | 0 |
| All TGAC v1 (500Kb) | 1 | 1 | 0.007 |

**Table 1D.**
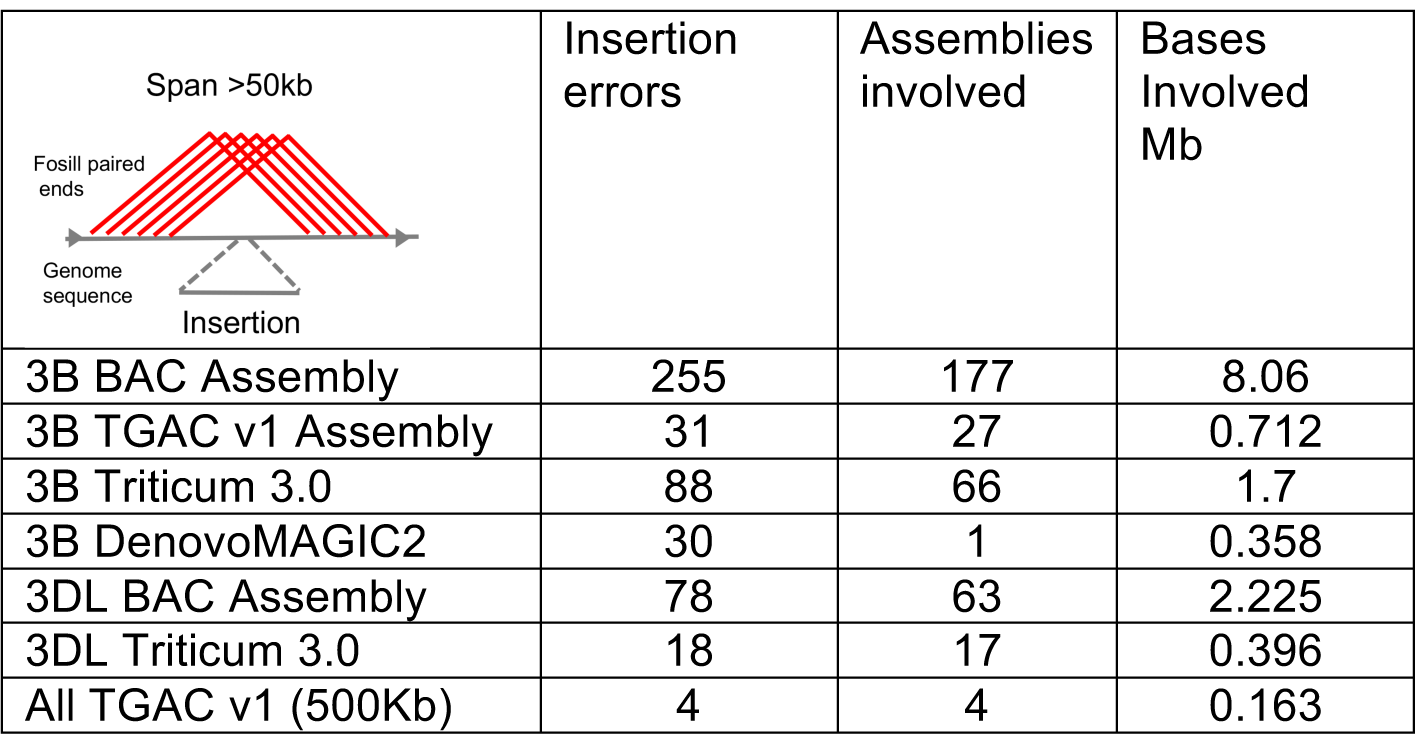
Potential erroneous insertions in scaffolds.

|  |  |  |  |
| --- | --- | --- | --- |
| Span >50kb | Insertion errors | Assemblies involved | Bases Involved Mb |
| 3B BAC Assembly | 255 | 177 | 8.06 |
| 3B TGAC v1 Assembly | 31 | 27 | 0.712 |
| 3B Triticum 3.0 | 88 | 66 | 1.7 |
| 3B DenovoMAGIC2 | 30 | 1 | 0.358 |
| 3DL BAC Assembly | 78 | 63 | 2.225 |
| 3DL Triticum 3.0 | 18 | 17 | 0.396 |
| All TGAC v1 (500Kb) | 4 | 4 | 0.163 |

**Table 1E.**
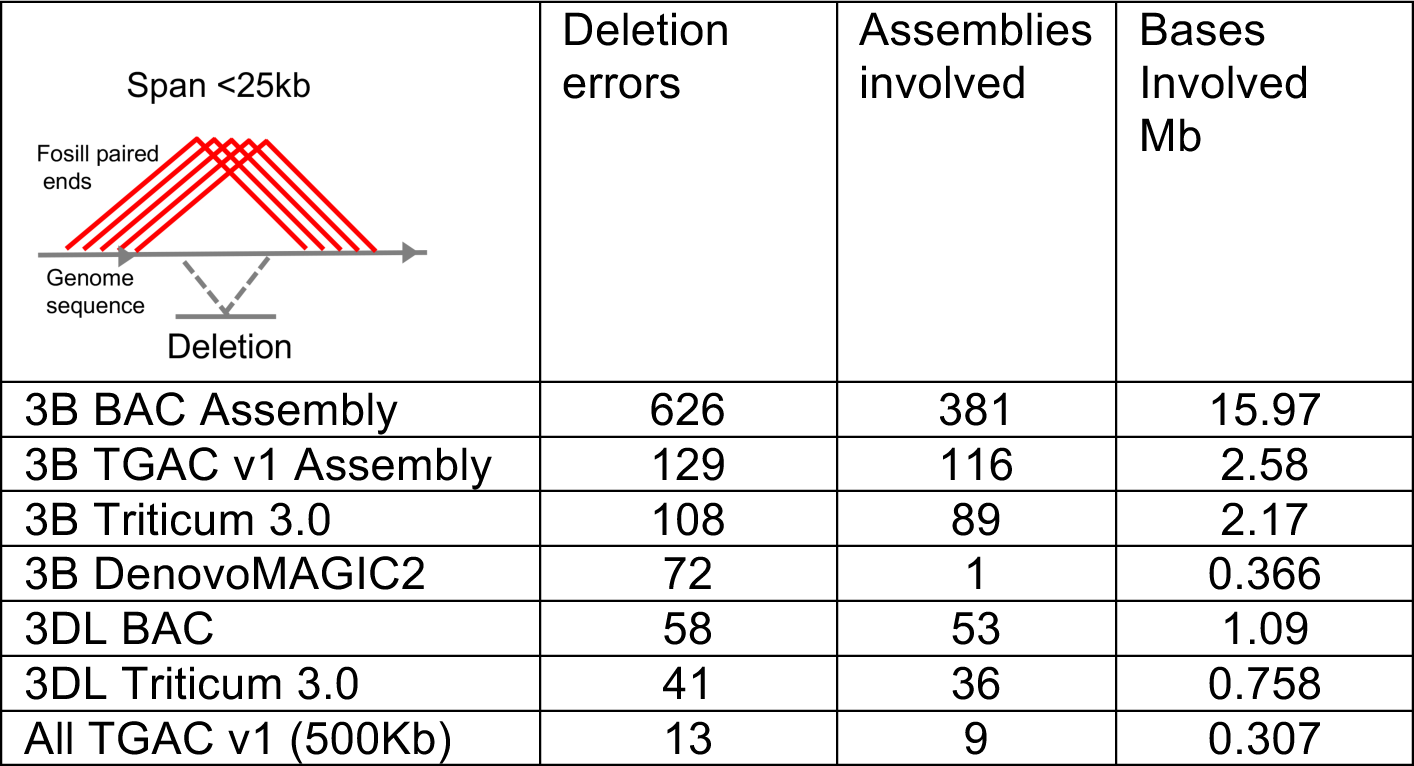
Potential erroneous deletions in scaffolds.

|  |  |  |  |
| --- | --- | --- | --- |
| Span <25kb<br>Fosmid paired ends<br>Genome sequence<br>Deletion | Deletion errors | Assemblies involved | Bases Involved Mb |
| 3B BAC Assembly | 626 | 381 | 15.97 |
| 3B TGAC v1 Assembly | 129 | 116 | 2.58 |
| 3B Triticum 3.0 | 108 | 89 | 2.17 |
| 3B DenovoMAGIC2 | 72 | 1 | 0.366 |
| 3DL BAC | 58 | 53 | 1.09 |
| 3DL Triticum 3.0 | 41 | 36 | 0.758 |
| All TGAC v1 (500Kb) | 13 | 9 | 0.307 |

Variation in Fosill insert sizes were consistent across the TGACv1, Triticum 3.0 and DeNovo Magic whole genome assemblies and the 3DL BAC assemblies. In contrast, chromosome 3B BAC assemblies had a higher variation of insert sizes (Table 1A). This may be due to a higher proportion of mis-assemblies in the 3B BAC assembly that could have introduced or removed small tracts of sequences, and possibly due to the use of a mixture of 454 and Illumina sequences. This variation in Fosill mate-pair matches did not contribute to assessment of assembly accuracy. The accuracy of assemblies was estimated by counting the bases included in correctly-sized windows (mean insert size */−sd) of Fosill mate-pair reads, and by the proportion of assemblies/scaffolds that were fully consistent with Fosill mate-pair windows along their length. The un-edited BAC-based scaffolds of chromosome 3DL were the least accurate, with only 17% of the assemblies covered with consistent fossil mate-pair matches, and 57% of the sequence included under consistent mate-pair matches (Table 1A). The 3B BAC assemblies, which have been extensively manually edited, were considerably more accurate, with 66% consistent assemblies and 85% of sequences in consistent windows. Looking at the TGACv1 3B assemblies, 61% of scaffolds were consistent and 80% of sequences were contained within consistent Fosill windows. In comparison, larger TGACv1 assemblies from the whole genome were all consistent with mate-pair windows and 99% of the sequences were in consistent windows. The differences with TGACv1 3B assemblies are most likely due to many shorter assemblies being included in the 3B assembly that limit the potential for 37 Kb mate-pair mapping, for example, there will be a low proportion of matches at the ends of assemblies. The Triticum 3.0 WGS assembly of 3B had 92% consistent assemblies, and 90% of sequences within consistent Fosill windows. Similarly, the Triticum 3.0 WGS assembly of chromosome 3DL had 99.5% assemblies and 80% of sequences in consistent windows. The DeNovo Magic WGS assembly of *T. turgidum* 3B contained 99.6% of sequences in consistent Fosill windows. As these assemblies were integrated into a single pseudomolecule the measure of the number of correct scaffolds was 100%.

Four different classes of discrepancies that may be due to assembly problems were assessed using Fosill mate pair mapping to assemblies: failed scaffolding, in which scaffolds had matches to only one end of Fosill end-sequences, and which may need to be broken; orientation errors in which the direction of one region of a scaffold is consistently reversed with respect to flanking regions; insertions, in which the span of Fosill mate-pair matches is greater than expected; and deletions, in which mate-pair spans are less than expected. These results are summarised in Tables 1B-1E. Of these potential error types, the most frequent were the potential erroneous joining of assemblies. These were highest in the BAC assemblies, and lowest in the DenovoMAGIC assembly of 3B. An example of this is shown in Figure 1B, where two BAC-based scaffolds were assembled at either end of chromosome 3B. Fosill mapping evidence, supported by TGACv1 assemblies, showed that the two scaffolds can be merged in opposite orientation to that originally assembled. Figure 2C reveals a 12 Kb deletion in a TGACv1 assembly that was due to a missing tandem duplication of the repeat, as validated by comparison with the Triticum 3.0 assembly. An aberrant insertion in a TGACv1 scaffold identified by Fosill mate-pair mapping was also validated by comparison with the Triticum 3.0 assembly (Figure 2D).

**Figure B.**
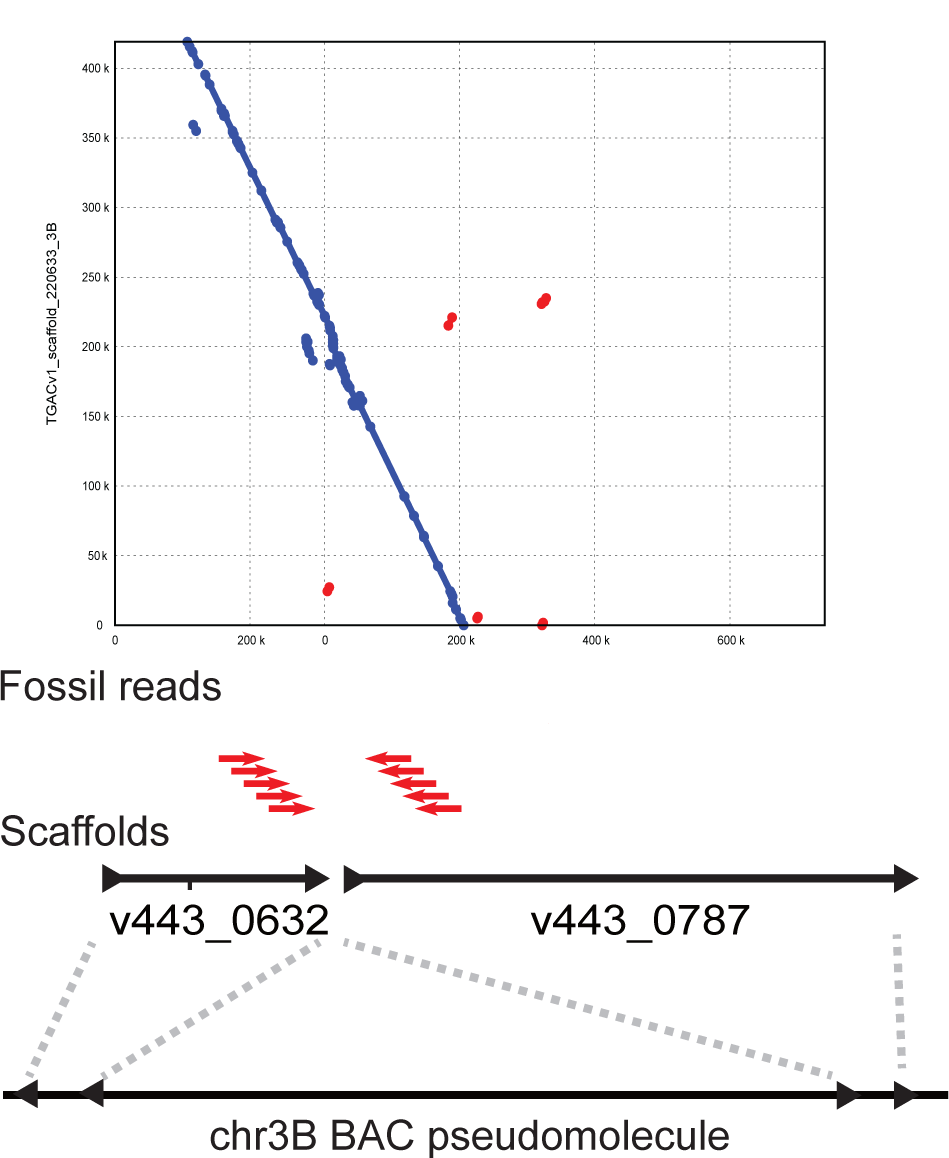
**B**. An example of a mis-join of the BAC-based assembly of chromosome 3B. Two scaffolds, v443_0362 and v443_0787, were originally assembled at opposite ends of chromosome 3B 730 Mb apart. Matches to Fosills indicated that these two scaffolds could be re-assembled together with v443_0362 in the opposite orientation. The Mummer plot shows that this join is supported by TGACv1 scaffold_220633_3B. Drawing not to scale.

**Figure C.**
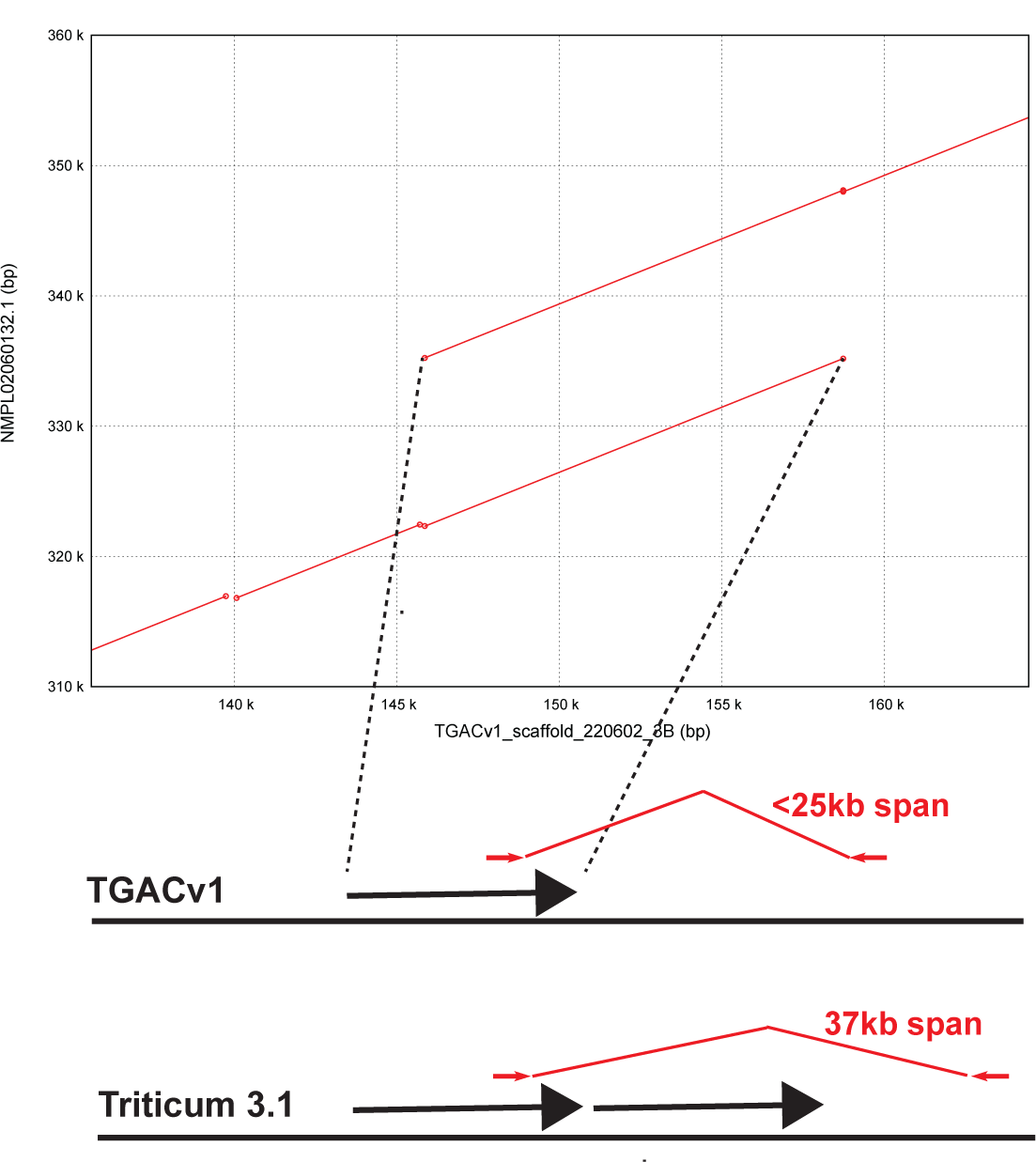
**C**. An example of an aberrant deletion in TGACv1 scaffold 220602 on chromosome 3B. Assembly missed a duplicate copy of a 12 Kb repeat (represented by an arrow) that was identified as a discrepancy in Fosill mate-pair matches. Comparison to a Triticum 3.0 scaffold identifies the predicted missing copy of the repeat. Drawing not to scale.

**Figure D.**
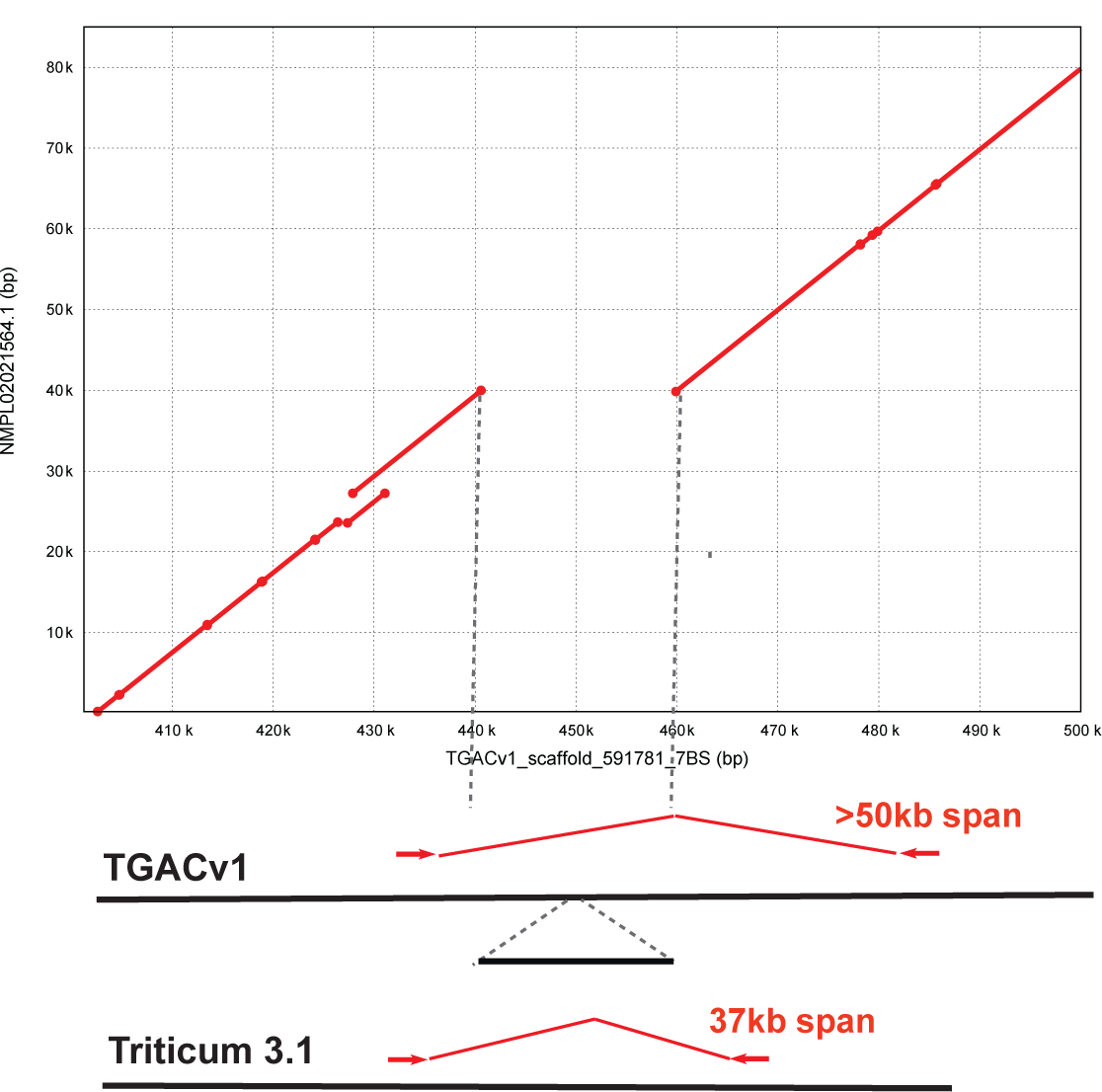
**D**. An example of an aberrant insertion in TGAv1 scaffold 591781 on chromosome 7BS detected by Fosill mate-pair matches of >50 Kb. Comparison to the Triticum 3.0 assembly of the same regions identifies the mis-assembled insertion. Drawing not to scale.

The TGACv1 large assemblies have relatively low numbers of mis-assemblies. The Triticum 3.0 assemblies of both 3B and 3DL had a consistently large number of potential mis-assemblies, with about 400-500 per chromosome or chromosome arm, affecting about 10 Mb of sequence region. Potential deletion errors, in which assemblies may be missing sequences, were most frequent in the BAC assembly of chromosome 3B, and were also the most frequent type of error in the DenovoMAGIC assembly. Deletions were least frequent in the TGACv1 whole genome assembly. Potential erroneous insertions were less frequent than deletions, with the highest rates of both types of potential error in BAC-based assemblies. In general, potentially erroneous deletions were more common in all assemblies than insertions. Mis-orientations were the rarest potential errors and were most prevalent in manual assembled 3B BAC scaffolds and essentially absent from TGACv1 and Triticum 3.0 assemblies, but were more frequent in the DeNovo Magic 3B assembly.

**Figure 3.**
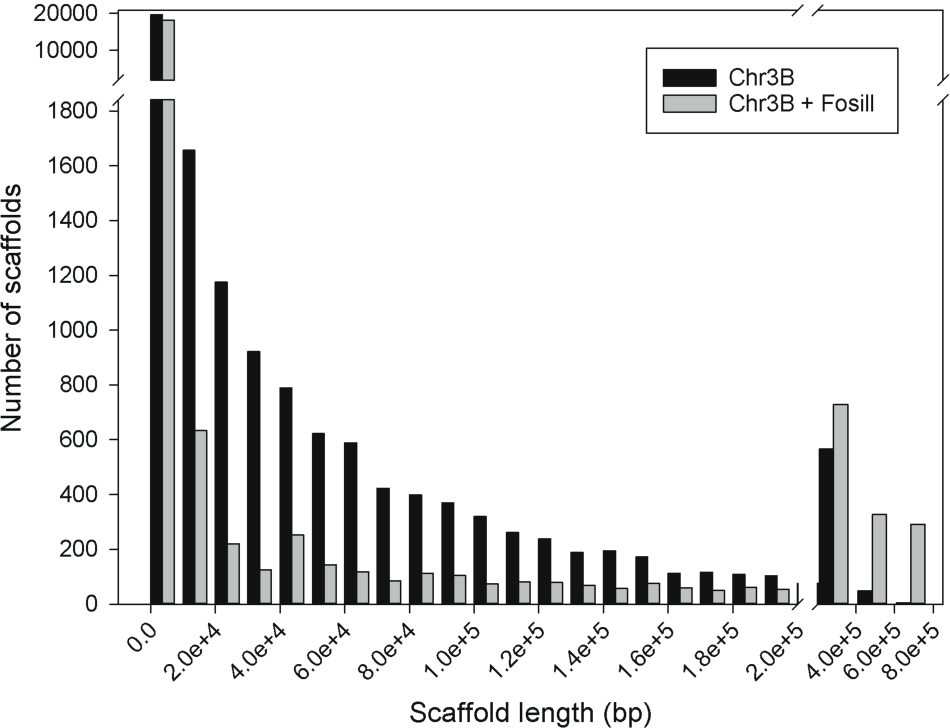
Increasing assembly contiguity using Fosill matches. **A**. Fosill mate-pair reads were used to link scaffolds of TGACv1 Illumina assemblies from chromosome 3B. The distribution of scaffold lengths and the number of scaffolds in each size range is shown before (dark bars) and after (grey bars) Fosill scaffolding. The numbers of smaller scaffolds are reduced, and the numbers of larger scaffolds are increased, by Fosill scaffolding, showing successful further assembly.

**Figure B.**
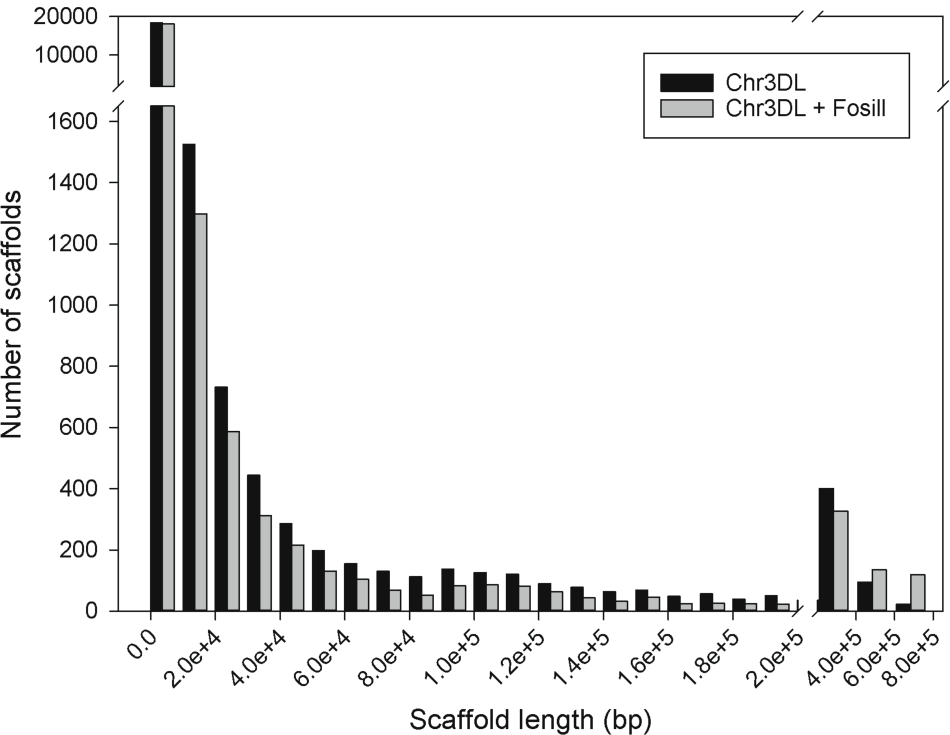
**B**. Fosill mate-pair reads were used to link scaffolds of BAC-based assemblies of chromosome 3DL. The distribution of scaffold lengths and the number of scaffolds in each size range is shown before (dark bars) and after (grey bars) Fosill scaffolding. The numbers of smaller scaffolds are reduced, and the numbers of larger scaffolds are increased, by Fosill scaffolding, showing successful further assembly.

### Using Fosill mate-pairs to create more contiguous assemblies

The wheat Fosill library was also used to create new joins in different assemblies. Table 2A shows that Fosill mate-pair reads made 267 new links between 477 chromosome 3B BAC scaffolds. Where available, TGACv1 3B assemblies spanning the new links precisely (124 cases), supporting the new join, and no examples were found where the new Fosill joins linked the wrong neighbours or the wrong strand. We then applied the Fosill mate pairs to make new joins in chromosome 3B TGACv1 assemblies and chromosome 3DL BAC assemblies. Table 2B shows the total assembly sizes were increased, while the number of scaffolds in the assemblies was decreased, and the scaffold n10 more than doubled in size. This showed, as predicted by simulations (Figure1), that 38 kb mate-pair reads can make new links that substantially improve contiguity of both WGS and BAC-based assemblies. Where available, independent assemblies supported these new Fosill-based links. Figure 3 shows the distribution of scaffold sizes and numbers before and after Fosill linking on TGACv1 chromosome 3B (panel A) and chromosome 3DL BAC (Panel B) assemblies. Increases in the numbers of larger assemblies and concomitant reduction in the numbers of smaller assemblies after Fosill joining was more apparent in the chromosome 3B WGS scaffolds than in the 3DL BAC scaffolds. This may reflect the fewer joins needed in the less fragmentary 3B assembly (2,808 scaffolds) that the very fragmented 3DL assembly (23,433 scaffolds).

**Table 2A.** Summary of new links made between BAC scaffolds on chromosome 3B.

| 267 new links | Strand | Validated by TGACv1 |
| --- | --- | --- |
| 37 links $\leq 40$kb on pseudomolecule | 20 correct strands | 12 |
|  | 17 reverse strands | 7 |
| 147 links $> 40$kb on pseudomolecule | 73 correct strands | 31 |
|  | 74 reverse strands | 38 |
| 83 links between scaffolds not assigned to the pseudomolecule | - | 36 |

**Table 2B.** Summary of changes in assemblies of chromosome 3B TGACv1 and chromosome 3DL BAC assemblies.

|  | **chr3B TGACv1** |  | **chr3DL BACs** |  |
| --- | --- | --- | --- | --- |
|  | **Before** | **After** | **Before** | **After** |
| Bases (bp) | 789,970,040 | 846,817,359 | 452,947,627 | 463,673,958 |
| Assemblies | 29,090 | 22,014 | 23,433 | 21,985 |
| Num>n50 | 2,020 | 644 | 790 | 721 |
| Min | 500 | 500 | 501 | 501 |
| n10 | 293,318 | 947,439 | 463,110 | 933,142 |
| n50 | 116,546 | 398,569 | 154,985 | 286,993 |
| Max | 739,616 | 2,867,878 | 1,240,092 | 1,942,124 |

**Figure 4.**
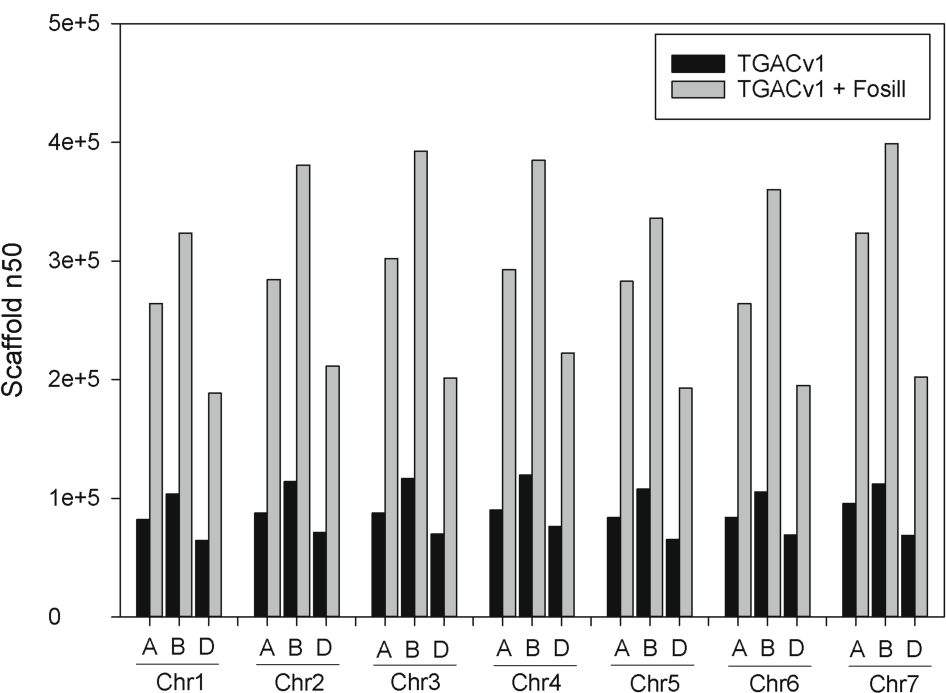
Fosill-mediated scaffolding of TGACv1 Illumina assemblies of the wheat genome. The 21 chromosomes are shown with their scaffold N50 values before (black bars) and after (grey bars) Fosill-mediated scaffolding.

Based on these improvements in both BAC-based and WGS scaffold contiguity by integrating Fosill mate-pair reads, we re-scaffolded the complete TGACv1 WGS assembly of the wheat variety Chinese Spring 42 [19]. Figire 4 and Supplemental File 1 show the scaffold sizes of each chromosome arm before and after integration of Fosill mate-pairs. Substantial increases in scaffold N50 of between 2.7 - 3.2-fold were achieved. The largest scaffolds increased is size between 1.5 - 3.2-fold, with the largest scaffold of 2.8 Mb on chromosome 3B.

## Discussion

Bread wheat is one of the three major cereals that we depend on for our nutrition, and generating accurate long-range assemblies is essential for new genomics-led approaches to crop improvement. However, its genome has been exceptionally challenging to sequence due to its polyploid composition of three closely-related large genomes, and extensive tracts of closely related repetitive sequences. Two strategies have been followed to deal with this genomic complexity: the first used BAC clones made from purified chromosomal DNA to reduce the complexity of chromosome-specific assemblies [27]; the second set of approaches uses different types of whole genome shotgun sequence technologies and assembly methods [19,20,22]. At this stage of wheat genome sequencing, when these complementary and contending approaches have been published, it is timely to assess the long-range accuracy of these different assemblies. For this, we mapped precise 38 Kb Fosill long mate pair reads to measure errors in different assemblies of chromosome 3B and the long arm of chromosome 3DL. We also used these Fosill mate pair reads to increase genome contiguity.

In order to maximise the accuracy of Fosill mate-pair read mapping to the A, B or D genomes and to repetitive regions of the hexaploid wheat genome, we modified the template conversion protocol of the Fosill 4 vector system [28] to generate longer paired 250 bp Illumina sequence reads. Nick-translation reactions to extend Nb.BbvCI nicks were optimised to generate an Illumina sequencing template between 750 - 1,000 bp. PCR amplification of re-circularised products was optimised to reduce amplification to the minimum required for efficient sequencing of a large library. Overall, 576.5M read pairs were generated from 55.1M clones (Additional File 2). When reads were mapped to chromosome 3B sequence assemblies [29], a consistent size distribution around 37.7 kb was observed (Figure 1A), demonstrating correct packaging and processing. Read depth varied several thousand-fold along chromosome 3B, likely due to matches of read-pairs to highly repetitive regions from across the genome. Consequently, only reads with depth ≤5 were used. Using this filter, we obtained sequence coverage of nearly 60% of the 833 Mb BAC-based chromosome 3B assembly. Simulations indicated that 0.75× sequence coverage of paired-end 250 bp reads was effective in creating long-range assemblies of wheat (Additional File 1), therefore we used Fosill read mapping for subsequent analyses.

Fosill reads were mapped to different assemblies of chromosome 3B and the long arm of chromosome 3D in order to compare the full range of current publicly available wheat genome assemblies. Several types of inconsistences spanning a wide range of scales have been detected by mapping long-range mate-pairs to human genome assemblies [24,30]. Tables 1A-1E) show the types of inconsistences detected in wheat assemblies using this approach. Looking first at the proportion of bases in different assemblies that were fully consistent with mapped 38 Kb mate-pair reads (Table 1A), the DenovoMAGIC Illumina-based assembly and the SMRT long-read Triticum 3.0 had respectively 99.6% and 90% of bases in consistent Fosill windows. The manually curated BAC-based assembly of 3B had 85% of consistent assembled sequences, while the TGACv1 3B assembly had 80% of assembled sequence in consistent windows, while the larger TGACv1 assemblies were 99% consistent. This difference may reflect the more fragmentary state of TGACv1 assemblies. The non-curated BAC assembly of chromosome 3DL was the least accurate according to this measure, with only 57% of bases in consistent windows. These data demonstrate both the superior accuracy of *de novo* whole genome sequencing strategies that incorporate deep and long 250 bp Illumina paired-end and mate-pair sequencing, and the relative accuracy of long-range assemblies generated by mate-pair assembly strategies [19,20,22], compared to BAC-based strategies ([27].

The most frequent type of inconsistency identified by Fosill mapping was the potential incorrect joining of assemblies (Table 1B). Illumina strategies produced the fewest incorrect joins, while BAC-based assemblies produced the most. Interestingly, assemblies of both 3B and 3DL made from PacBio SMRT reads combined with 150 bp Illumina paired end reads (forming mega-reads) [22] had more possible assembly issues than Illumina-only assemblies. While a more complete assembly and relatively long assemblies were achieved from SMRT sequences, these assembly issues suggest that reads longer that 10 kb, or including long Illumina mate-pair libraries, could further improve SMRT-based assemblies. Assembly methods may also need further optimisation to utilize fully the potential of SMRT long reads. Furthermore, integrating long 250 bp Illumina reads into mega-reads may improve assemblies by distinguishing very closely related sequences, such as repeat regions from homoeologous chromosomes.

Potential deletion events were also quite common in all assemblies, and were the most common inconsistencies detected in DenovoMAGIC assemblies of 3B. The sizes of these events are not known precisely, but they have a minimum size of 12 Kb (Table 1E). These probably arise from missing tracts of near-identical sequence in assemblies. Similarly, potential insertions may arise from the incorrect integration of near-identical sequences into assemblies. The observation that potential deletions are more frequent than potential insertions suggests that all WGS-alone assembly strategies could achieve more complete assemblies of the wheat genome such as that achieved using PacBio SMRT sequence assemblies. Finally, potential mis-orientations/inversions of assemblies are more common in the DevoMAGIC assembly of 3B that the other whole-genome assemblies. Although this approach has yet to be fully described, mis-orientations may reflect more relaxed criteria for linking scaffolds than related Illumina-based assembly and scaffolding approaches ([19].

How much more accurate can the best current assemblies of bread wheat and wild emmer wheat be, judging by their assemblies of chromosome 3B? Fosmid end-mapping to 2005 versions of human genome assemblies [24] identified 297 longer range discrepancies in the 3.2 Gb genome. Scaling from chromosome 3B (0.8 Gb) with 127 potential inconsistences, our analyses predict 480 discrepancies per 3Gb of wild emmer wheat genome assembly-roughly twice the error frequency of 2005 versions of the human genome. It is highly likely that a DenovoMAGIC version of the hexaploid bread wheat genome will achieve similar high levels of accuracy and coverage.

Three-fold increases in the scaffold N50 sizes of the TGACv1 whole genome assembly were achieved by an additional scaffolding step using Fosill mate-pairs. In addition to making a more useful genomic resource, this additional scaffolding shows the relatively fragmentary but highly accurate TGACv1 assembly has the potential for substantial further improvement. Considering the urgent need to generate accurate long-range assemblies of multiple elite bread wheat genomes and wild progenitor species for crop improvement programmes, linked read technologies [31] and Nanopore long reads [32] provide promising new opportunities for efficient, cost-effective and open-source approaches to identifying of a wide range of structural and phased sequence variation in wheat genome assemblies.

## Methods

Detailed descriptions of experimental and computational procedures are shown in Additional Files. These describe simulation of 38 Kb mate-pair reads for assembly (Additional File 1), Production and sequencing of Fosill libraries (Additional File 2) and physical mapping and sequencing of BACs from chromosome 3DL (Additional File 3).

### General bioinformatics

All analytical pipelines have been deposited in GitHub, and relevant links are shown in the manuscript and Additional Files. Joinable read pairs from Illumina Miseq or HiSeq sequencing were removed using FLASH v1.2.11 [33]. Ligation adaptors in reads were trimmed off using CutAdapt v1.6 [34]. Sequencing primer sequences and low-quality sequences in reads were removed using Trimmomatic v0.32 (parameter) [35]. Then the resulting reads were evaluated using FastQC v1.2.11 [36].

Trimmed reads were further filtered using ReadCleaner4Scaffolding pipeline https://github.com/lufuhao/ReadCleaner4Scaffolding). Both mates in each pair was mapped to chr3B BAC scaffolds using bowtie v1.0.1 [37]. And then the Picard MarkDuplicates (v1.108, http://broadinstitute.github.io/picard) was used to remove the duplicates as single reads. A read depth threshold was used to remove the repeat-like reads by plotting the summary of output from samtools depth, and the reads mapped to those regions with higher depth were not used for scaffolding. The remaining reads were subjected to removal again as pairs. Those reads mapped to multiple positions, whose mates were not mapped, or had the wrong orientation, were removed. A window size filter was applied to identify sets of ≥5 neighbouring reads in sliding windows of less than 10 Kb that had all their mates in a following window of less than 20 kb. Variations of the expected distance between mate-pairs (average ± sd) of approximately 3 sd was used to identify potential assembly discrepancies.

## Data Availability

Fosill mate-pair reads from Chinese Spring 42 in this study have been submitted to the EBI European Nucleotide Archive (ENA), and are available in study accession PRJEB23322. Chromosome 3DL BAC scaffolds are available in ENA study accession PRJEB23358.

## Declarations

The authors declare they have no competing interests.

## Authors’ contributions

MWB conceived and coordinated the project, and wrote the manuscript. F-HL planned and carried out bioinformatics analyses, NMcK constructed the Fosill libraries, GK and MDC sequenced chromosome 3DL BACs and managed sequence data, and DH managed all sequencing library production and Illumina sequencing.

## Acknowledgements

We are grateful to Louise Williams (Broad Institute) for Fosill vectors and detailed advice.

## Additional File 1

### Genome assembly simulation

Before adopting Fosill jumping libraries for analysing and improving wheat genome assemblies, we simulated assembly processes using Fosill mate-pair reads on three of the largest scaffolds of the BAC-based assembly of chromosome 3B [1]. Simulations used Next-Generation illumina SIMulation PipeLinE (NGSimple, https://github.com/lufuhao/NGSimple) to assess various parameters, including library types, fragment sizes, read length, and sequencing depth. Simulated reads were generated by the Mason program [2] and then trimmed by Trimmomatic version 0.32 [3]. Quality control was applied to these datasets before and after, in order to confirm the removal of the low quality and low complexity bases in reads. Velvet v1.2.10 [4] was used to assemble these reads from different parameter settings in one run. And finally, MUMmer v3.23 [5] was used to map the assembled contigs back to its original scaffold to evaluate the quality of the faux assemblies. Results for a single representative scaffold are shown below.

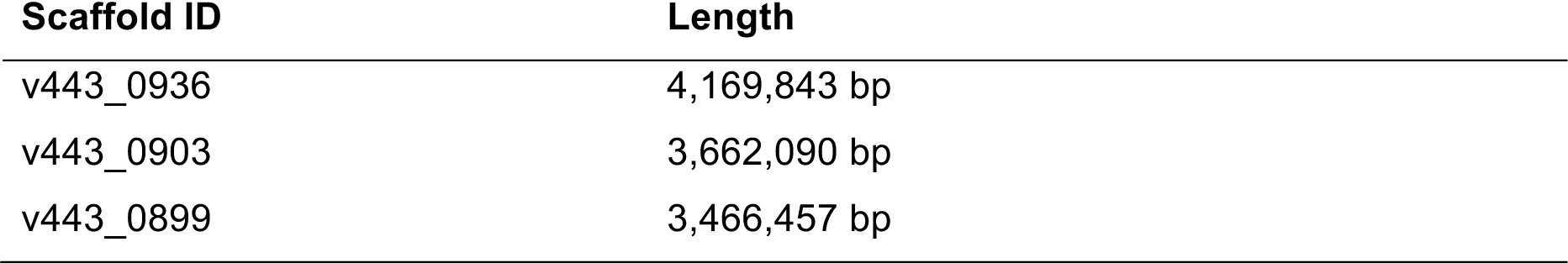
Chromosome 3B scaffolds used for simulation

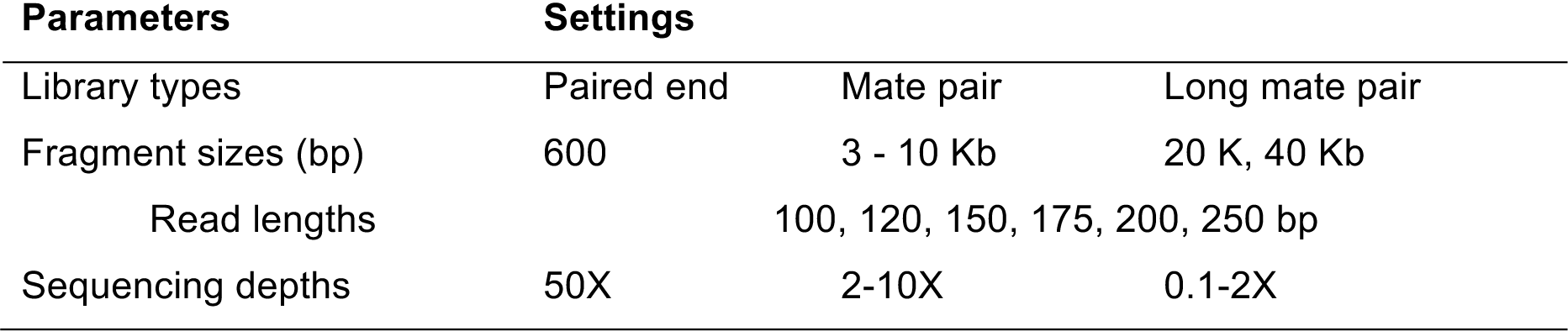
Simulation parameter settings

#### 1.1 Optimizing mate-pair fragment length

Simulation settings for mate-pair library fragment sizes

1.1 Optimizing mate-pair fragment length Simulation settings for mate-pair library fragment sizes
|  | Insert size | Coverage | Read length |
| --- | --- | --- | --- |
| Paired-end | 600 bp | 50x | 250 bp |
| Mate pair | 3,000-10,000 bp | 10x | 250 bp |
| Long mate pair | 20 Kb, 40 Kb | 2x | 250 bp |

**Fig. 1.**
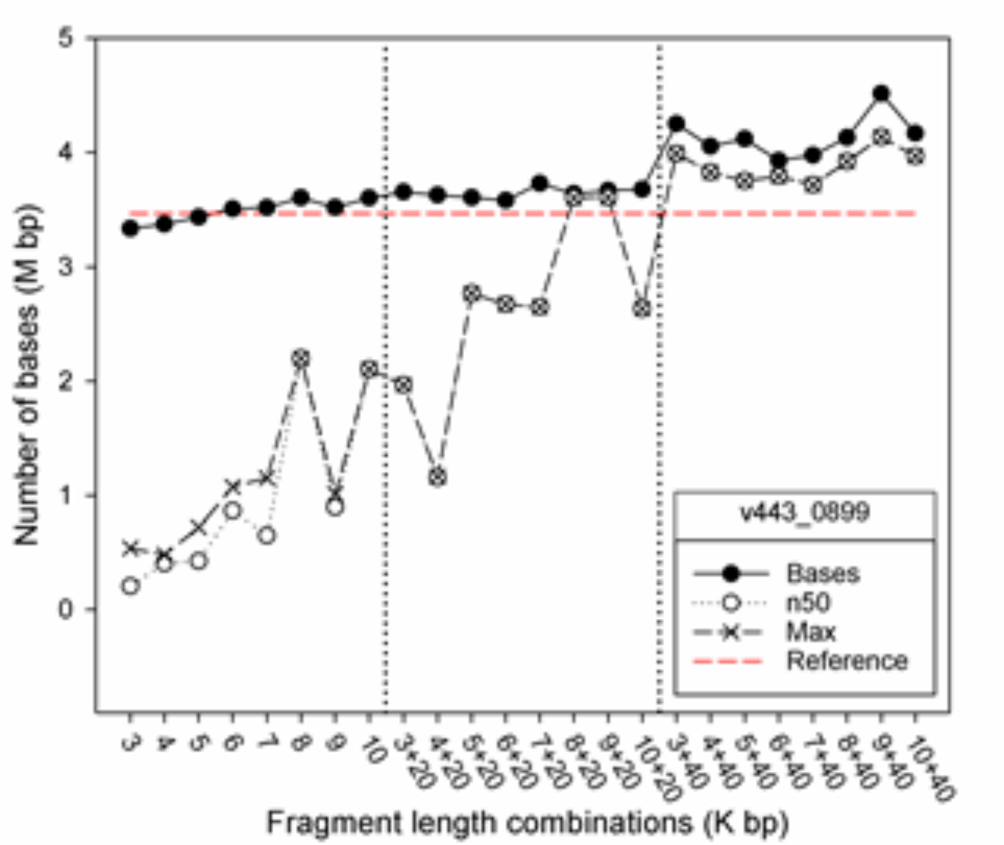
Summary of the *de novo* assemblies based on scaffold v443_0899

**Fig. 2.**
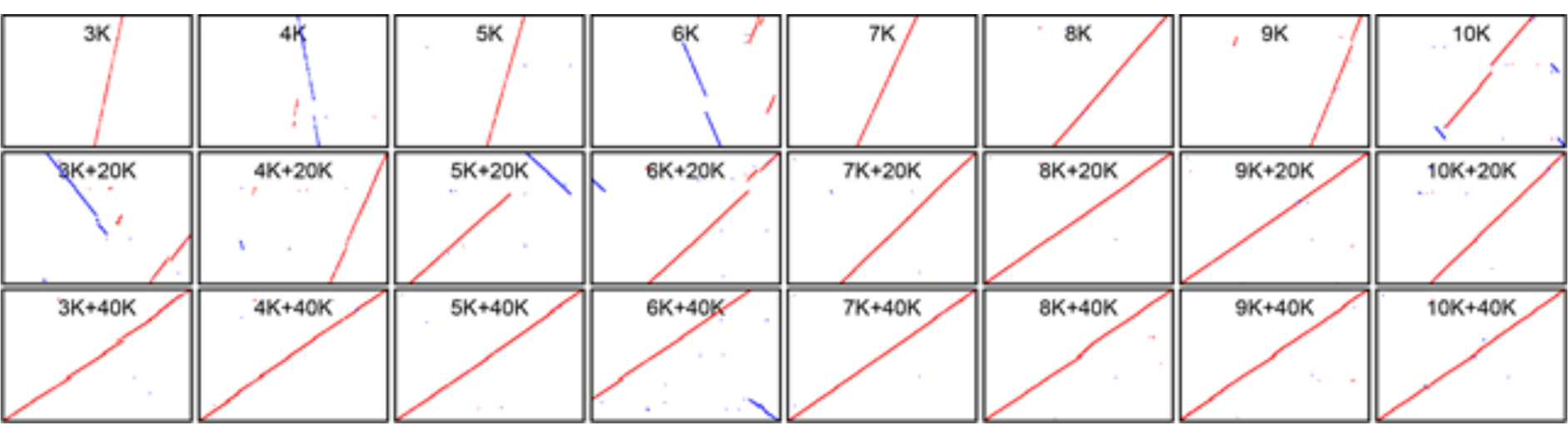
Plot of longest contig against its reference v443_0899

#### 1.2 Optimizing mate-pair sequence read length

**Table 2.** Simulation settings for sequence read length

|  | Insert size bp | Coverage | Read length |
| --- | --- | --- | --- |
| Paired-end | 600 | 30x | 100-300 bp |
| Mate paired | 7,000 | 10x | 100-300 bp |
| Long mate paired | 40,000 | 2x | 100-300 bp |

**Fig. 3.**
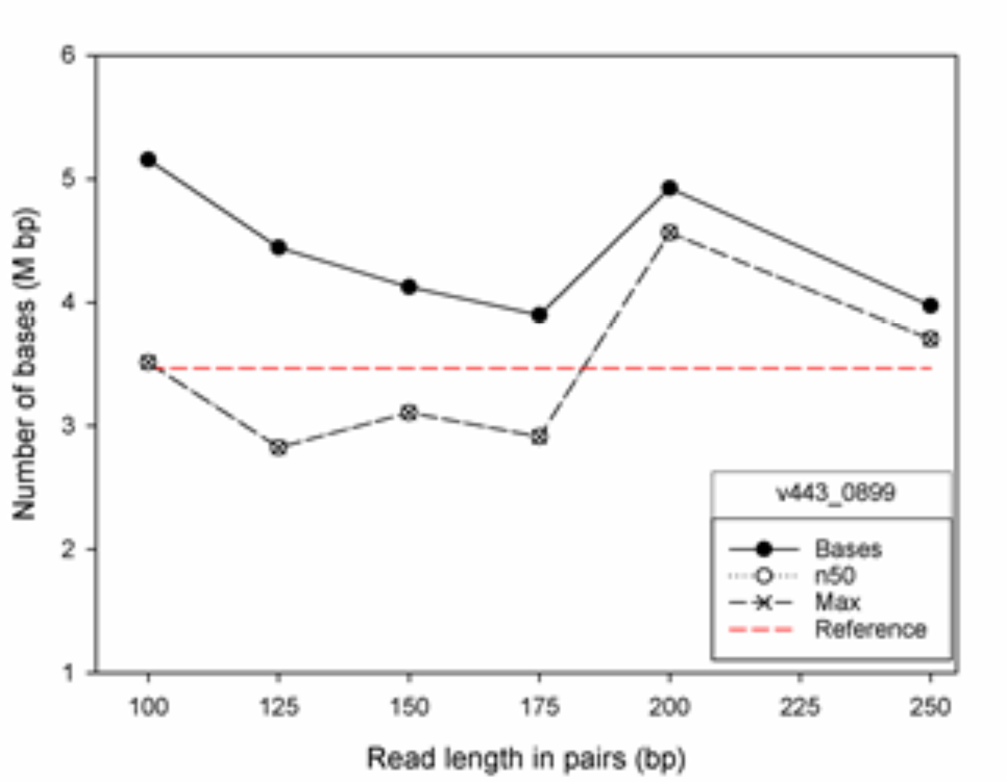
Summary of the *de novo* assemblies based on v443_0899

**Fig. 4.**
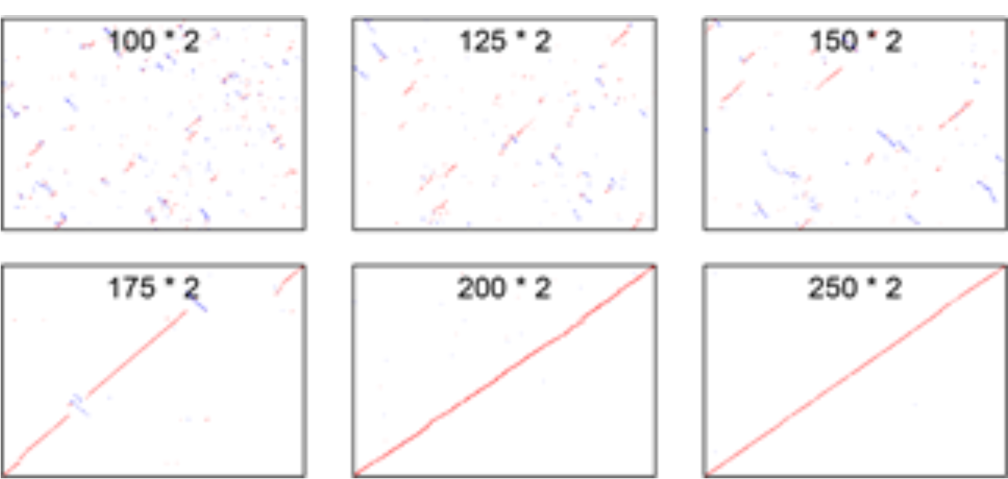
Plot of longest contig against its reference v443_0899

#### 1.3 Optimizing long mate-pair coverage

**Table 3.** Simulation Setting for coverage by long mate-pair libraries

|  | Insert size bp | Coverage | Read length |
| --- | --- | --- | --- |
| Paired-end | 600 | 50x | 250 bp |
| Mate paired | 7,000 | 10x | 250 bp |
| Long mate paired | 40,000 | 0-2x | 250 bp |

**Fig. 5.**
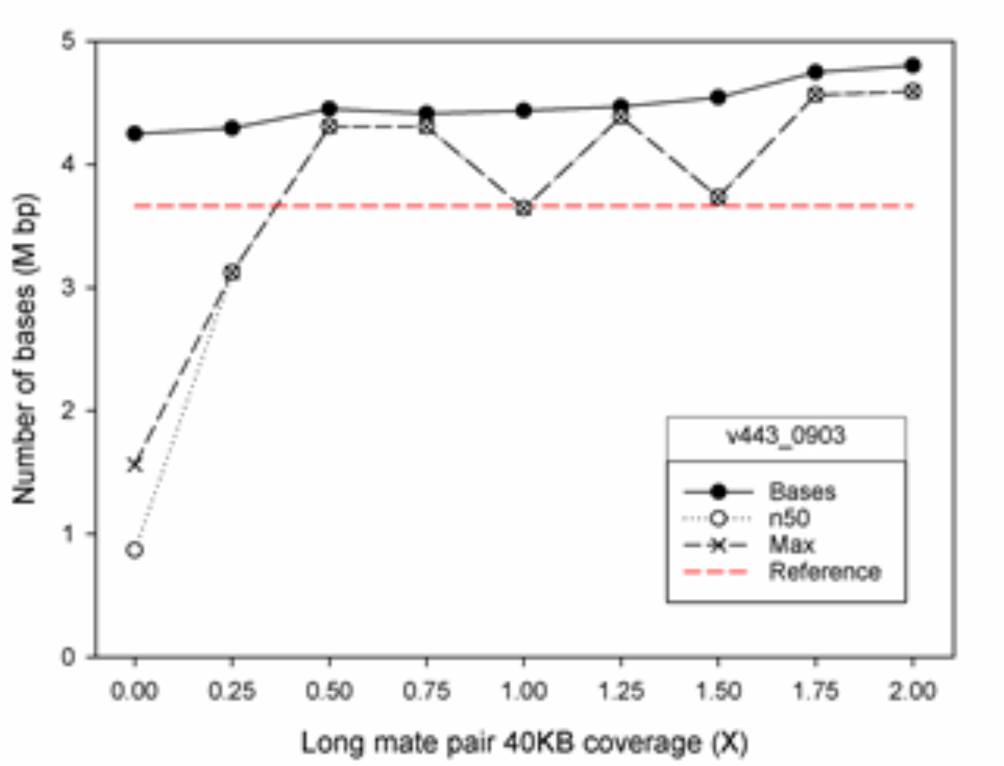
Summary of the *de novo* assemblies based on v443_0903

**Fig. 6.**
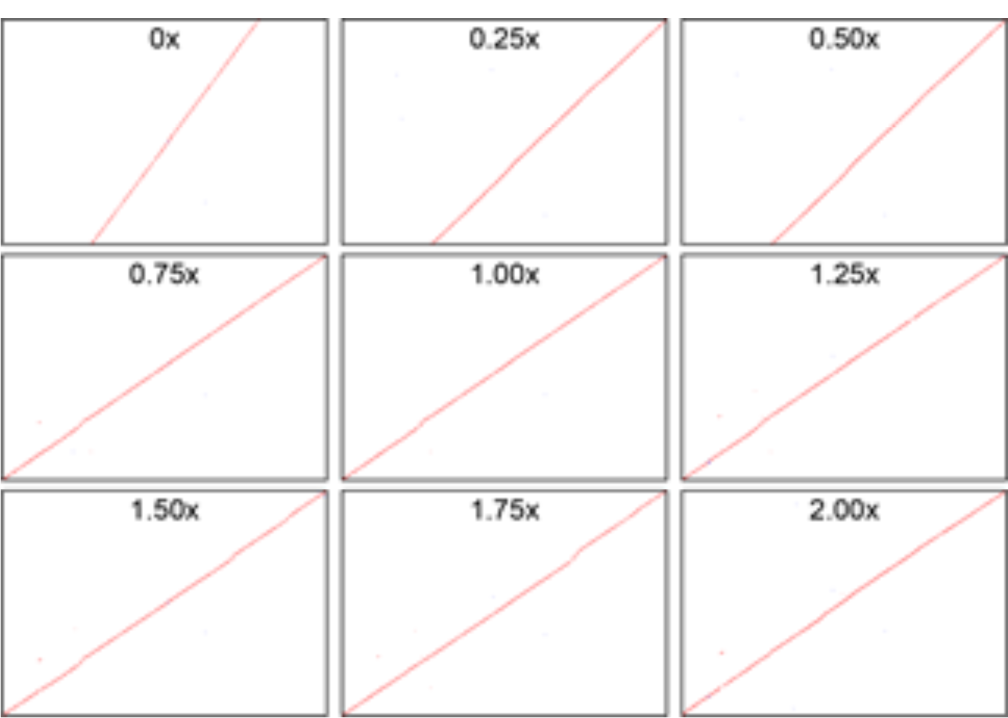
Plot of longest contig against its reference v443_0903

**Table 4.** Simulation Setting for coverage by mate-pair libraries

|  | Insert size bp | Coverage | Read length |
| --- | --- | --- | --- |
| Paired-end | 600 | 50x | 250 bp |
| Mate paired | 7,000 | 2-10x | 250 bp |
| Long mate paired | 40,000 | 1x | 250 bp |

**Fig. 7.**
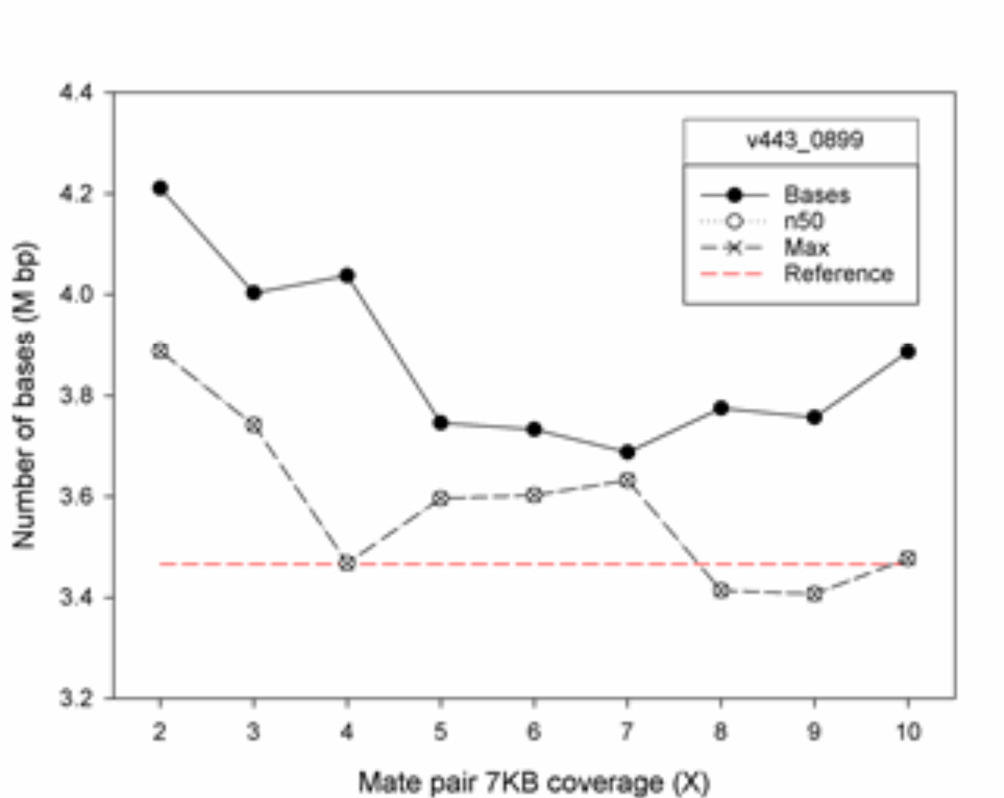
Summary of the *de novo* assemblies based on v443_0899

**Fig. 8.**
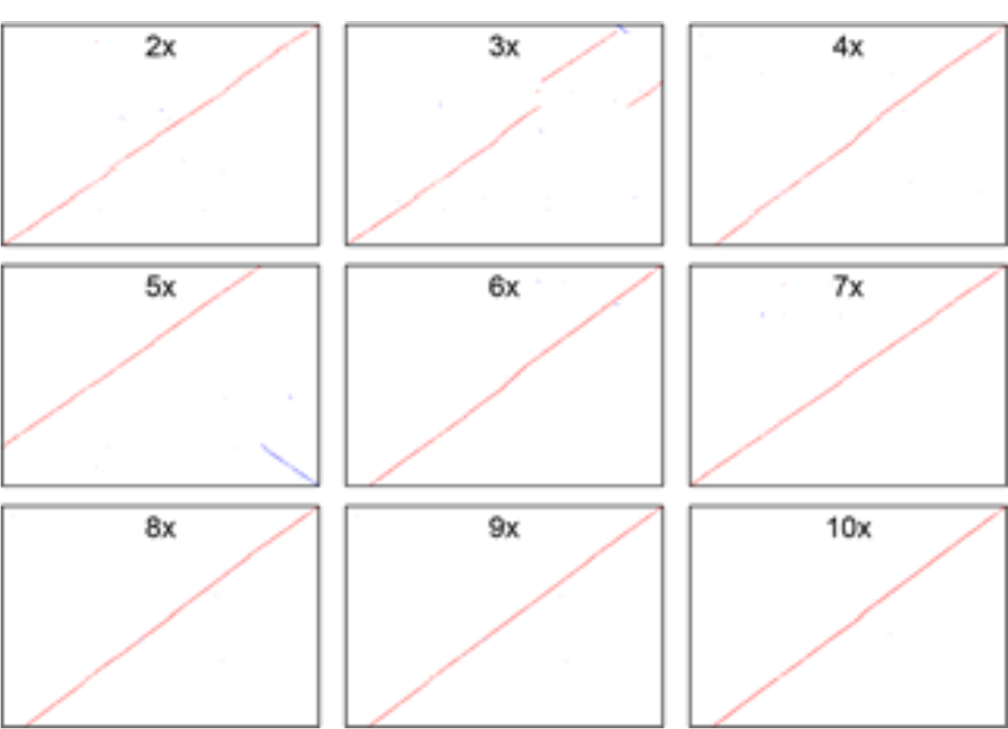
Plot of longest contig against its reference v443_0899

## Additional File 2

### Fosill library production

The pFosill 4 cloning vector was used in this study was kindly provided by Louise Williams (Broad Institute, Cambridge, MA, USA). The construction, preparation and downstream steps for non-size-selected DNA fragments were carried out as described [1]. Only differences in the methods are described here.

A single-seed-descent line of *Triticum aestivum* Chinese Spring (CS42) was used for high molecular weight DNA extraction as described [2]. 30 μg of Genomic DNA was sheared to approximately 40 Kb average fragment size by HydroShear (Digilab, Marlborough, MA, USA) using the Large Shearing Assembly set at speed code 40 for 20 cycles. Sheared DNA was assessed by Pulse Field Gel Electrophoresis. 0.5-1 μg of sheared and non-sheared DNA was run on BioRad CHEF DRII. 120 degrees, 6V/cm, 1 to 10 second ramp switch, 17 hours at 14 °C. DNA was visualised by staining gel with Ethidium Bromide (Figure 1).

10 μg batches of sheared DNA was end-repaired in 175 μl reactions containing 1 X T4 ligase buffer, 0.25 μM dNTPs, 15 units T4 DNA polymerase, 50 units T4 polynucleotide Kinase, and 5 units Klenow fragment (all NEB) for 30 mins at 20 °C. TE was added to the DNA up to 400 μl, which was then cleaned and concentrated through Amicon Ultra 0.5 ml 100k concentrator (Millipore) at 2,000 g to approximately 30 μl. Recovered DNA was measured on the Qubit fluorometer (Invitrogen) using Quant-iT dsDNA BR kit.

t-b index linker A: GATCTCTACCAGG and t-b index linker B: CCTGGTAGAG were annealed and multiple ligations were set up using 500 ng end repaired genomic DNA and 200-fold molar excess of annealed linker. DNA was pooled and between 5 (500 μl) to 20 (2000μl) ligations were cleaned and concentrated with one Amicon column.

Multiple 10μl ligations were set up containing 250 ng linkered DNA and 500ng cut and dephosphorylated pFossill 4 vector. 10μl ligation was packaged with 2 successive 50 μl MaxPlax λ packaging extract (Epicentre) for 90 mins at 30 °C. 1,850μl Phage dilution buffer and 140 μl DMSO were added (2,100 μl total volume) The libraries were titered and stored at −80°c. A -competent GC10 (Sigma) was used for processing packaged libraries into fosmid DNA. A proportion of 1.5 ml λ packaged sample to 40 ml of cells was found to give best transformation efficiency. Cultures were grown overnight at 30 °C in LB.

Fosmid DNA was isolated from LB culture using Qiagen’s Plamid Maxi Purification Kit. 20 ml of Solutions P1, P2 and P3 were used and the supernatant was transferred to a new tube through a layer of Miracloth prior to addition to the Maxi column. Fosmid DNA was eluted with 500 μl TE and quantified using a Qubit fluorometer.

### Conversation of Fosmids into Fossills

Pools of approximately 2.5 million independent Fosill clones were collected (see Table 1) and 10 μg of DNA from each pool was processed, with the following modifications. 900 ng was nicked with Nb.BbvCI for between 55-60 mins, before S1 nuclease treatment. 300 ng of DNA was re-circularised in 650 μl containing 1× T4 ligase buffer and 8,000 units of T4 ligase (NEB) at 16 °C for 16 hours. Products were purified using a Qiagen PCR cleanup kit. Columns were washed twice with 750 μl wash buffer and eluted with 55 μl TE.

A trial PCR was used to determine minimal amplification required for Illumina template preparation. 2 μl of re-circularised DNA was amplified in 25 μl total volume of 1x Phusion HF master mix and 0.5 μM PCR primers:

SBS3: 5’AATGATACGGCGACCACCGAGATCTACACTCTTTTCCCTACACGACGC 3’ SBS12:

5’AAGCAGAAGACGGCATACGAGATGATCGATCGTGACTGGAGTTCAGACGTGTGC 3’ Cycling parameters were 98 0C for 3 mins, 16 and 18 cycles respectively of 98 0C for 15 secs, 65 °C for 30 secs, 72 °C for 30 secs and a final extension at 72 °C for 7 mins. PCR products were analysed with a MultiNA Bioanalyser (Shimadzu) using DNA 12000 reagent kit in on-chip mode. An intensity measurement of 5-8 MV, which equated to approximately 7.5 ng/μl to 12ng/μl for the 700 - 950 bp peak, was optimal. Following analysis of MultiNA data to determine minimal cycling conditions for each pool, Super-Pools of approximately 10 million independent Fosill clones were selected from the pools and minimal cycle number calculated for each pool to give sufficient material for sequencing. A total of 24 50 μl PCR reactions each containing 4 μl of Fossill DNA for each Super-Pool. Cycling parameters, primers and primer concentration were same as for trial PCR (except for varied cycle numbers). PCR products from Super-Pools were combined (1,200 μl) and purified with AMPure XP beads and eluted with 40 μl of TE. 4 μl of sample was used for MultiNA analysis to confirm size range and quantity. 30 μl of sample was size-selected on 1.5% agarose cassette with R2 marker using Sage Science BluePippin (Beverly, MA, USA) set to collect fragments between 650 - 1,000bp. Successful size selection was confirmed using TapeStation size measurement, and DNA was purified with AMPure XP beads and eluted in 25 μl TE. Sequencing was performed using 2 × 250 bp pair-end sequencing chemistry on an Illumina HiSeq 2500 sequencer.

**Figure 1.**
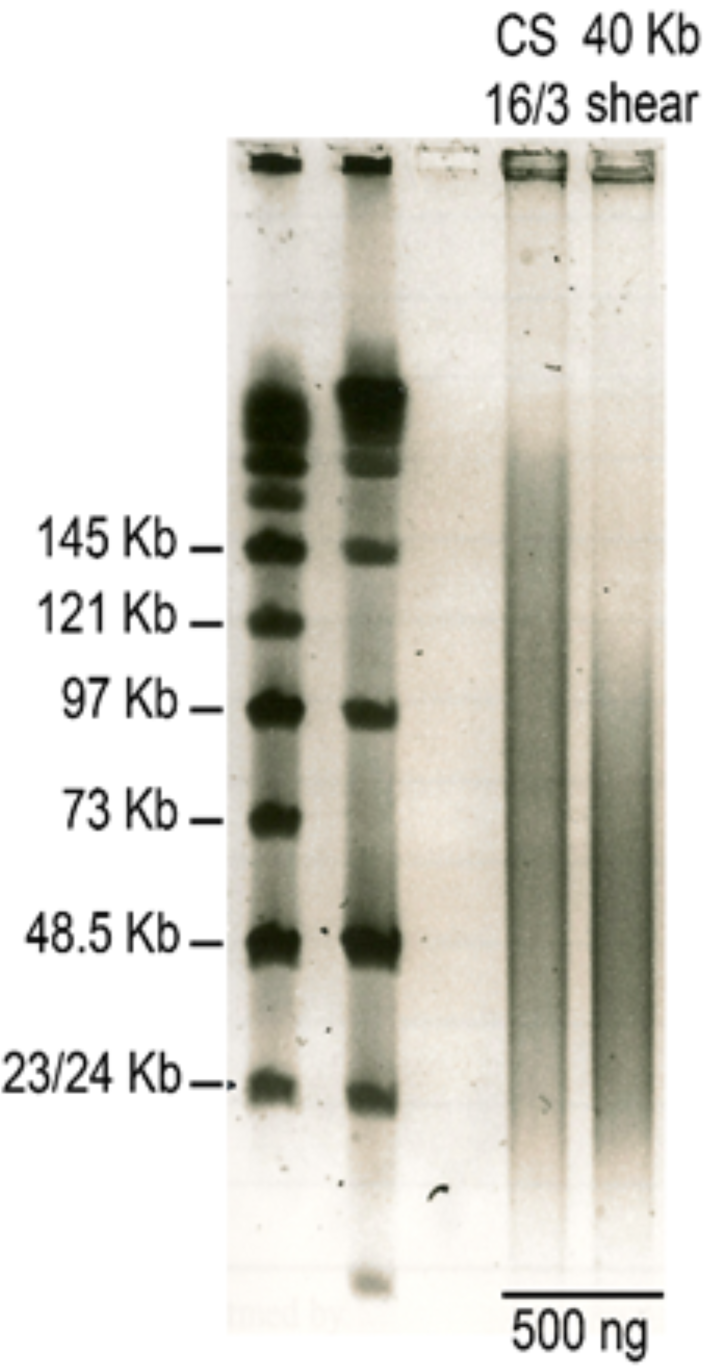
PFGE analysis of sheared DNA for Fosill vector cloning.

**Table 1.** Summary of Fosill libraries and paired-end reads generated.

| Library | Titre (M) | Platform | Num_Read | PCR redundancy |
| --- | --- | --- | --- | --- |
| Lib17562 | 0.90 | MiSeq | 7,283,029 | 6.05 |
| Lib18185 | 5.50 | HiSeq | 51,668,481 | 9.74 |
| Lib18186 | 5.80 | HiSeq | 55,092,287 | 9.35 |
| Lib19454 | 10.09 | HiSeq | 124,755,368 | 11.68 |
| Lib19455 | 9.99 | HiSeq | 117,879,434 | 9.14 |
| Lib19456 | 11.59 | HiSeq | 111,113,269 | 6.94 |
| Lib19457 | 11.64 | HiSeq | 108,714,323 | 6.89 |
| Total | 55.51 | - | 576,506,191 | 8.51 average |

## Additional File 3

### Fosill mate-pair mapping

Read pairs from Illumina Miseq or HiSeq sequencing that joined were removed using FLASH v1.2.11 [1]. Ligation adaptors and vector sequences in reads were trimmed off using CutAdapt v1.6 [2]. Sequencing primer sequences and low-quality sequences in reads were removed using Trimmomatic v0.32 (parameter) [3]. Resulting reads were then evaluated using FastQC v1.2.11 [4].

Trimmed Fosill mate-pair reads were filtered using the ReadCleaner4Scaffolding pipeline https://github.com/lufuhao/ReadCleaner4Scaffolding. Both mates of each pair were mapped to chr3B BAC scaffolds using bowtie v1.0.1 [5]. Picard MarkDuplicates (v1.108, http://broadinstitute.github.io/picard) was then used to remove duplicates as single reads. A read depth threshold was determined to remove any highly repetitive reads by plotting the summary of output from samtools depth, and all the reads mapped to those regions with depth >5 were not considered for scaffolding. Reads mapping to multiple positions, whose mates were not mapped, or were in the wrong orientation, were removed. A window sizing method was used to map mate-pairs to genomic regions. A group of ≥5 neighbouring mate reads within a “driver” window of less than 10 Kb were linked by the average 37.7 Kb insertion size °/− sd to a “follower” window of 20 Kb. Figure 1 shows that nearly all mate-pairs mapped to chromosome 3B using these criteria. Reads mapping within these window criteria were used to define regions of chromosomes that were consistent with the average insertion size, or had inconsistent matches.

To generate coordinates of each scaffold on the 3B pseudomolecule, BAC scaffolds were mapped to the pseudomolecule and plotted using SyntenyDraw (available on https://github.com/lufuhao/SvntenyPlot). These coordinates were compared with our evidence from ReadCleaner4Scaffolding pipeline. To validate mapping, mate-pairs were mapped TGAC v1 chr3B contigs.

Filtered Fosill mate-pair reads were mapped to the TGACv1 whole genome assembly of Chinese Spring 42 as described above, using SSPACE v3.0 to join scaffolds with five fossils links.

**Figure 1.**
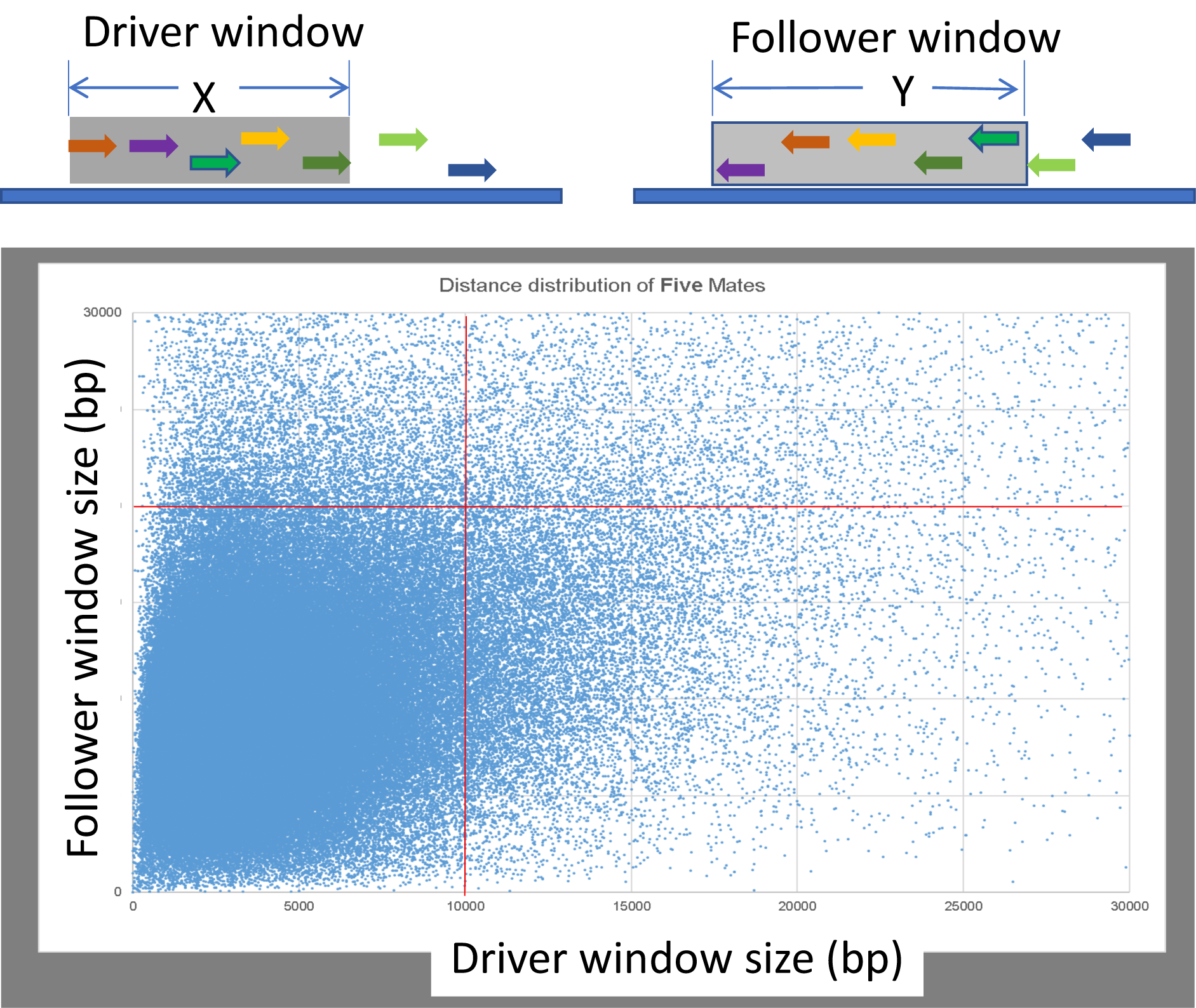
**Mapping Fosill mate pairs to genomic scaffolds**

The distribution of mate-pair links of five adjacent Fosill mate-pairs in different sized windows to their corresponding mate-pair read at the average insert size °/− sd was mapped on chromosome 3B BACs. The vast majority of mate-pairs in a 10 Kb window were found in 20 Kb windows at the correct distance. These window sizes were used to map Fosill mate-pairs to genomic scaffolds.

## Additional File 4

### Sequencing Chromosome 3DL BAC minimal tiling path

#### BAC library preparation and sequencing

The 3DL BAC library was prepared from flow sorted chromosomes [1] at The Institute of Experimental Botany, Olomouc, Czech Republic, and was fingerprinted at CNRGV (Toulouse, France) using SNaPshot-based high information content methods [2]. The raw fingerprint data was processed according to IWGSC guidelines. LTC [3] was used to build the physical map and generate a minimum tiling path (MTP). The final path consisted of 620 fingerprint contigs (FPCs) containing 5 or more BACs. 6,338 BACs were selected for sequencing (6,252 MTP clones plus 86 bridge clones).

Paired-end and long mate-pair (LMP) libraries were prepared and sequenced to generate PE reads for each BAC and a pool of LMP reads for each 384 well plate of BACs. LMP reads were processed as described in [4]. After standard QC, filtering and de-multiplexing, the reads were ready for assembly.

#### BAC assembly and mate-pair preparation

Reads were aligned to *E. coli* DH10B, wheat chloroplast and mitochondria sequences using Bowtie 2 [5]. Read pairs with one or more reads mapping with 95% identity or above were removed. Reads were also aligned to the pIndigoBAC-5 BAC vector sequence. Read pairs where one or more reads mapped to the middle of the vector sequence were removed while pairs where a read mapped to the end of a vector sequence were kept. This identified vector insert ligation sites. BACs were assembled individually using ABySS [6]. The BAC assemblies had an average insert size of 112,886 bp and an average N50 of 25,214 bp. In addition to the pooled LMPs, we used the 9 Kb and 12 Kb whole genome wheat LMPs from [4]. These were first filtered for non-3DL reads by alignment to the IWGSC CSS assembly where all 3DL contigs were replaced with our BAC assemblies. Reads were assigned to individual BACs as a side effect of this process.

Reads were then assigned from the pooled LMPs to each BAC. A Jellyfish [7] 31-mer hash table was generated from each assembly and these were combined to create a table of 31-mers found in the BACs on each plate. To identify LMP reads matching BACs, the “sect” function of the Kmer Analysis Toolkit (KAT) [8] v1.0.5 was used to generate a ft-mer coverage profile of each LMP read in each pool using plate-specific PE ft-mer hash tables. The plate-specific LMP reads were then classified to individual BACs on that plate using ft-mers from individual BAC assemblies.

#### Chromosome arm assembly

Before any scaffolding, the BAC assemblies belonging to each FPC were then merged to remove redundancy. This was done first using CD-HIT [9] and then BLAST [10]. Any overlapping sequence at the end of two BAC contigs of at least 98 % identity and 1000 bp in length resulted in the two contigs being merged into one new contig. Following this procedure each FPC had an average size of 460 Kb and an N50 of 17 Kb.

The non-redundant FPCs were then scaffolded using Soapdenovo [11] with the assigned pooled and whole genome LMP reads. This resulted in an average FPC size of 782 Kb and an N50 of 180 Kb. Finally, the FPC sequences were combined and the merging process was run again. This resulted in a total size of 455 Mb for the whole chromosome arm and an N50 of 145 Kb.

